# Prior information differentially affects discrimination decisions and subjective confidence reports

**DOI:** 10.1101/2022.10.26.513829

**Authors:** Marika Constant, Michael Pereira, Nathan Faivre, Elisa Filevich

## Abstract

According to Bayesian models, both decisions and confidence are based on the same precision-weighted integration of prior expectations (“priors”) and incoming information (“likelihoods”). This assumes that priors are integrated optimally and equally in decisions and confidence, which has not been tested. In two experiments, we quantitatively assessed how priors inform both decisions and confidence. With a gamified dual-decision task we controlled the strength of priors and likelihoods to create pairs of conditions that were matched in posterior information, but differed on whether the prior or likelihood was more informative. We found that priors were underweighted in discrimination decisions, but used to a greater extent in confidence about those decisions, and this was not due to differences in processing time. With a Bayesian model we quantified the weighting parameters for the prior at both levels, and confirmed that priors are more optimally used in explicit confidence, even when underused in decisions.

---

Human perception has often been shown to be based on a Bayesian inference process, in which the brain infers information about the environment by integrating incoming information with previous beliefs^1^. Computationally, this involves the integration of a prior distribution (“prior”) with a likelihood distribution (“likelihood”) to give a posterior distribution (“posterior”), which then forms the basis of a belief or percept. Many studies have shown evidence supporting the idea that perception and decision-making can be explained as Bayesian inference^1–7^. Further, confidence, i.e. the sense of certainty that typically accompanies perceptual decisions, can also be explained by Bayesian inference models.

In formal terms, Bayesian models propose that confidence corresponds to the perceived posterior probability of being correct about our inferences, based on the relative strengths of the posterior probabilities of each hypothesis being considered. This Bayesian confidence model has been tested and supported empirically, in both animals and humans^8–14^. For example, evidence supporting the Bayesian confidence model has been found at the neural level in rats, with firing rates producing a signature “folded-X” pattern^10^. In humans, this same Bayesian confidence model can quantitatively capture confidence reports, as well as predicting critical qualitative signatures of confidence, including the same characteristic folded-X pattern^14^. The Bayesian approach has been highly influential (although non-Bayesian alternatives have been proposed^15,16^). Importantly, this framework considers that the posterior percept is based on the precision-weighted integration of priors and likelihoods, and also that this same posterior gives rise to decisions and confidence. This simple formulation relies on two assumptions: first, that both decisions and confidence integrate priors and likelihoods optimally; and further, that confidence integrates priors and likelihoods in the same way as decisions. But these assumptions must be empirically tested, as the alternative is also possible: either confidence, decisions, or both, might integrate priors in systematically biased ways, and might do so asymmetrically.

Meanwhile, there is evidence to suggest that these kinds of suboptimalities often occur in human perception and confidence. Various systematic biases have been found in the information that enters confidence, such as a bias towards decision-congruent evidence^17^, or an overweighting of perceived sensory noise^18,19^. Furthermore, several studies have found that confidence incorporates different or additional information compared to decisions^20–25^. This supports the possibility that there are asymmetries in the way that certain sources of information influence these different processing levels. With regard to prior information, empirical work examining Bayesian confidence models has typically used uninformative, ‘flat’ priors, so it has not been possible to detect these potential biases or asymmetries. Two recent studies have begun examining confidence under informative priors, and found that confidence thresholds liberalise following prior-congruent stimuli^26,27^. These results show that priors influence confidence, but still cannot answer how optimal this influence is, and how it compares to the influence on decisions. In order to understand how our sense of confidence arises across different situations in which we may have highly informative prior expectations, as well as to rigorously test Bayesian confidence models, it is critical to understand quantitatively how priors are weighted relative to likelihoods in confidence computations, and how this relates to their weighting in decisions. More broadly, due to the pervasive role of priors in our processing, studies assessing the Bayesian confidence model under informative priors are important for generalizability.

Here, we examined (1) whether the use of prior information is optimal relative to the use of new sensory information, and (2) regardless of optimality, whether prior information is used the same way at the level of decisions and confidence. We did this both behaviourally and by fitting a generative Bayesian model with free weighting parameters that allowed us to quantify the relative use of priors at the level of decisions and confidence. Additionally, we assessed whether any possible asymmetries in the use of the prior at these different processing levels could be explained by differences in evidence accumulation time.

In two experiments, participants completed a gamified dual-decision task in which they made right/left decisions about two consecutive dot motion stimuli per trial. Critically, participants were informed about the added rule that, following correct responses to the first (‘lead’) stimulus, the second (‘target’) stimulus would go to the right. Conversely, incorrect decisions about the lead stimulus would be followed by leftwards-moving target stimuli. This meant that, in an optimal Bayesian observer, the prior expectation for a rightward target stimulus would be equal to the decision confidence about the lead stimulus. In other words, if participants were very confident that their response to the lead stimulus was correct, they would have a very strong prior that the target would be a rightward stimulus. Conversely, if participants were very unsure about their response to the lead, they would have equal expectations that the target would go right or leftwards. A dual-decision task with this same rule was used recently^28^ to investigate the influence of the prior on the decision level, which was interpreted as a measure of implicit confidence. Here we build on that work in order to assess the potentially differential role of priors in decisions and in explicit, subjective confidence ratings.

In our design, we also made use of the task structure in order to be able to vary the strength of priors and likelihoods on the same dimension. Only this way could we directly, behaviourally compare the relative influence of priors and likelihoods on responses. We did this by building two conditions that were matched in the amount of total available posterior information, but differed in whether the lead or target was more informative. This allowed us to measure whether accuracy, confidence, and metacognitive efficiency differed between these two conditions, which could indicate either over- or underweighting of the priors relative to likelihoods. Going further, in order to quantify that weighting in both decisions and confidence, and test the precise way in which the use of prior information might differ at these different processing levels, we fit a Bayesian model to the data with parameters capturing the weighting of the prior in decisions and confidence. In the second experiment, we then further investigated whether potential asymmetries between the weighting of priors in decisions versus confidence could be simply attributed to differences in processing time.

## Results

### Experiment 1

#### Dual-Decision Task and Conditions

On each trial of the gamified dual-decision task, participants (N=21) saw two consecutive random dot motion stimuli and made a decision after each about whether the coherent motion was to the right or left (Figure 1A), followed finally by a confidence rating about the second decision. We told participants that their task was to herd a flock of sheep (represented as dots in the RDK stimulus) towards the barn on the right of the screen. We gamified the rule (linking correct lead-decision responses to rightwards-moving target stimuli) and asked participants to position a sheepdog by responding to the lead stimulus. If the sheepdog was in the correct place, the sheep (coherently moving dots) of the target stimulus would go to the right. This meant that participants’ internal decision confidence about the lead stimulus formed the strength of their prior for a rightward target stimulus. In this way, the strength of the prior could be manipulated by changing the coherence of the lead stimulus (with L: low, M: medium, or H: high coherence), and the strength of the likelihood could be manipulated by changing the coherence of the target stimulus (also L, M, or H). This allowed us to create two conditions with the same available posterior information to the optimal observer (‘posterior level’): One in which there was more prior information due to a stronger lead stimulus (Stronger-Lead) and one in which there was more new sensory information due to a stronger target stimulus (Stronger-Target) (Figure 1B). These two conditions existed in matched pairs across three overall levels of available posterior information (L_post_: L+M or M+L, M_post_: L+H or H+L, and H_post_: M+H or M+H). With this experimental design, we were able to assess whether participants’ accuracy, confidence and metacognitive performance depended on the condition.

**Figure 1.**
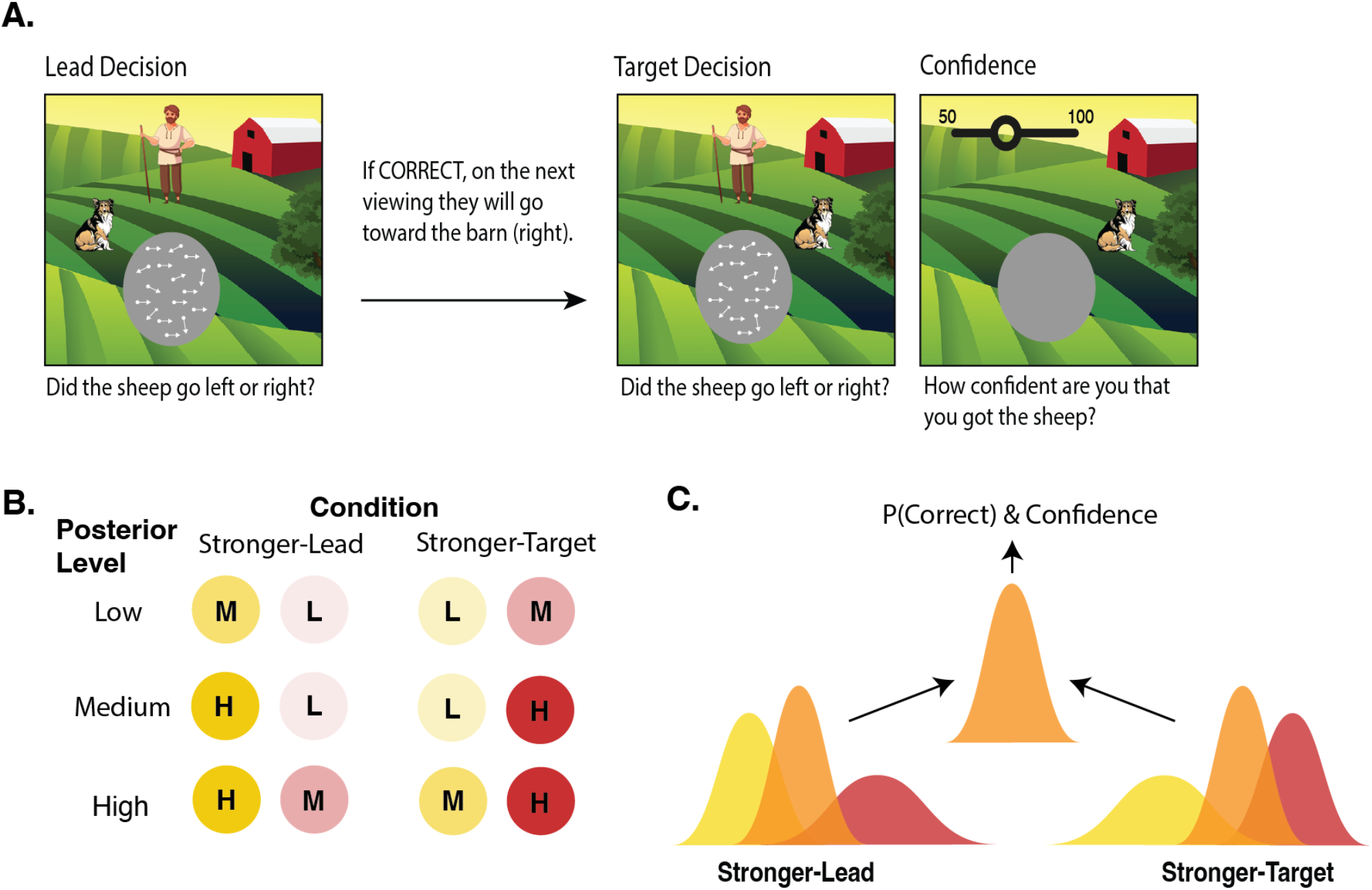
Task and Conditions Sketch. **A. Gamified dual-decision paradigm**. On each trial, participants viewed and made right/left decisions about two consecutive dot motion stimuli (lead and target), which they were told represented flocks of sheep. We explicitly informed participants that if they were correct about the first decision, then the target stimulus would be going to the right, and if incorrect, then it would be going to the left. This meant that, in an optimal observer, the prior for a rightward target stimulus should be equal to the lead decision certainty. They also rated their confidence in the target decision. In Experiment 2, the paradigm was the same except there was a 2-second delay after viewing the target stimulus, before participants were allowed to make the target decision. **B. Conditions**. We manipulated the coherence of the lead and target stimuli (each of which could have L: low, M: medium, or H: high coherence), here depicted with the circle transparency, to create two conditions that had matched available posterior information but differed in whether the lead or target stimulus was stronger. This was tested at three overall posterior levels - low posterior information (L+M), medium posterior information (L+H), and high posterior information (M+H). **C. Sketch of the Stronger-Lead vs Stronger-Target Manipulation**. The posterior percept (orange) should optimally be the precision-weighted integration of the prior (yellow), which in our task was always rightward and could span from 50-100%, and likelihood (red). Both conditions led to the same available amount of posterior information, which (in the optimal case) leads to the same probability of a correct choice as well as confidence. Hence, the target accuracy and confidence will only differ between conditions if the two sources of information are not integrated optimally in the decisions and/or confidence. Note that this is only a sketch aimed at conveying the intuition of how the conditions were matched in terms of posterior information. The prior for a rightward target stimulus is captured more accurately by a step function, shown in Figure 7.

#### Manipulation Check

We first ensured that response accuracy increased with increasing coherence of the stimuli, indicating that we effectively manipulated internal signal strength as intended. Additionally, we investigated whether participants used the task structure as we wanted them to, using the rule and hence their prior to guide their target decisions, at least to some extent. If this were true, we expected participants to perform better on the target decisions of each trial, on which they had additional information from the prior (lead stimulus) to guide their choice. To test both these predictions we built a logistic regression model on response accuracy with fixed effects of information level (L, M, H), decision order (lead or target), and their interactions (see Table 1 for the model syntax). We found a significant interaction between decision order and information level (*χ*^2^(2)=43.53, p<0.001, BF_10_ = 9.71 × 10^4^), with differences between information levels reduced in the target decisions. However, post-hoc pairwise comparisons revealed that response accuracy remained significantly higher with increasing information levels for both decisions (all p<0.001), suggesting our coherence manipulation to work as planned (Figure 2A). Additionally, response accuracy was higher in the target decision compared to the lead decision at each information level, and this was significant at the low and medium information levels (both p<0.001), revealing appropriate use of the task structure.

**Table 1.**
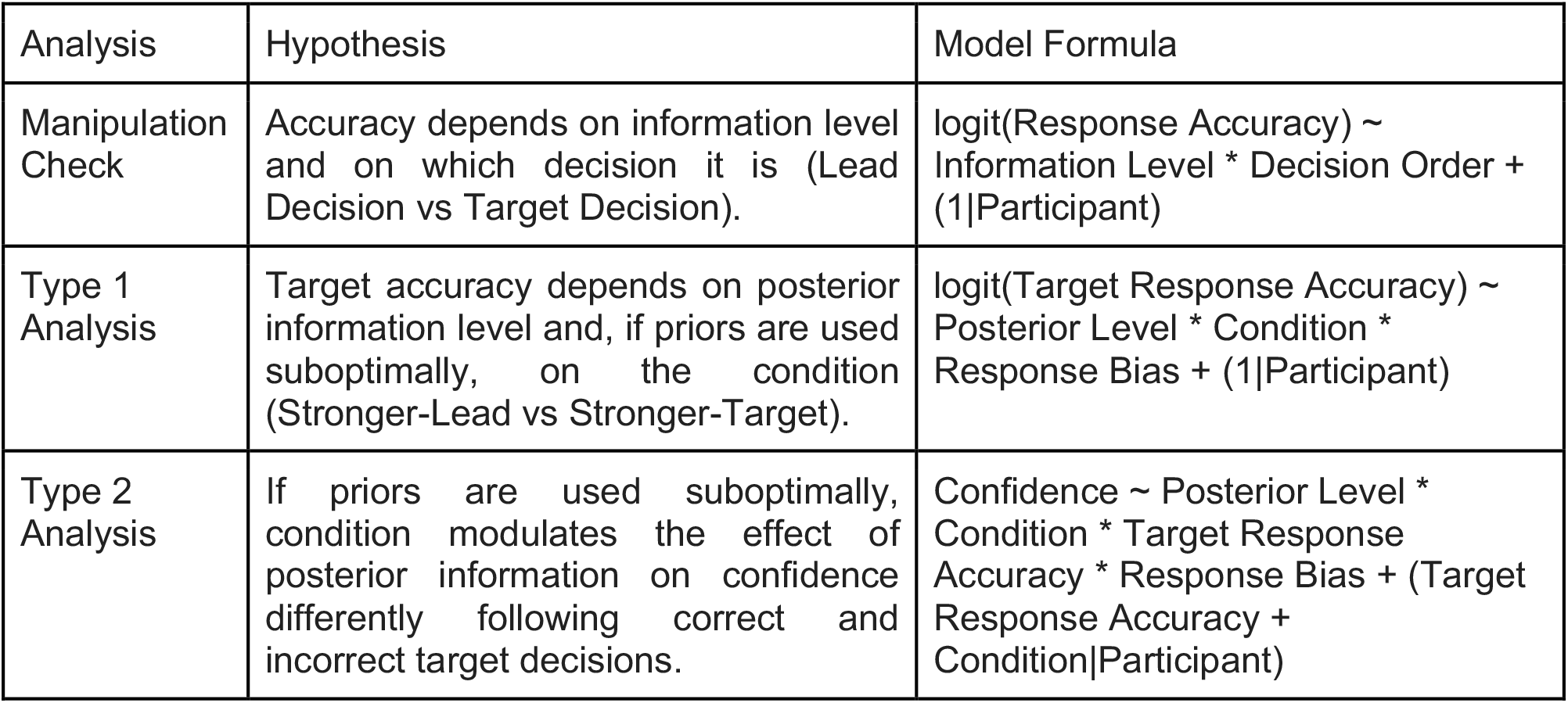
Regression Models

**Figure 2.**
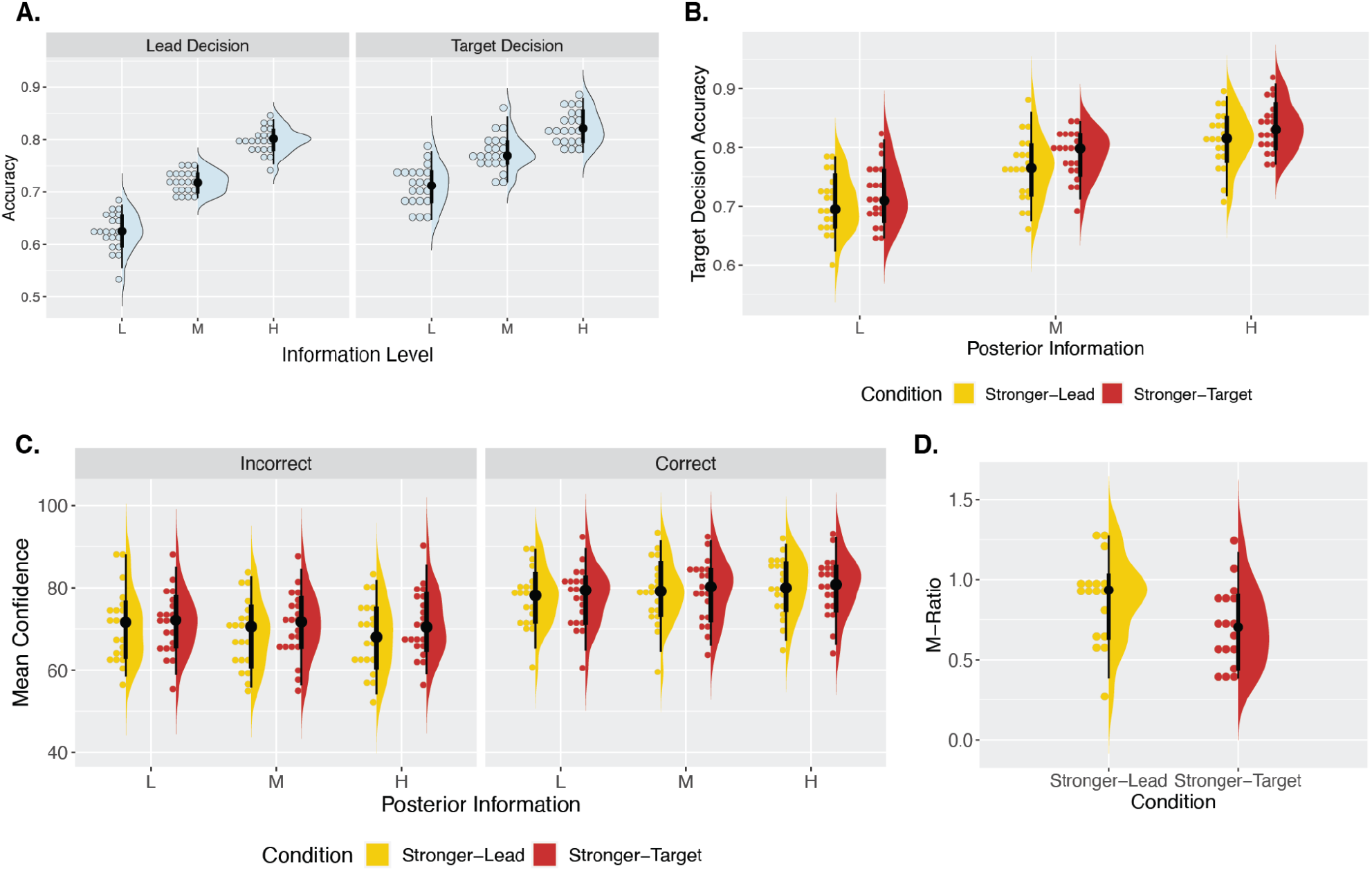
Behavioural Results. **A. Manipulation check**. As expected by experimental design, increasing the information level increased accuracy in both the lead decisions (M_L_Lead_=0.63, SD=0.04; M_M_Lead_=0.71, SD=0.02; M_H_Lead_=0.80, SD=0.02) and target decisions (M_L_Target_=0.71, SD=0.04; M_M_Target_=0.78, SD=0.03; M_H_Target_=0.82, SD=0.03), suggesting that changes in dot coherence successfully increased the strength of the internal signal. Accuracy was also higher in the target decisions compared to the lead decisions, suggesting at least some use of the task structure and prior information. In the raincloud plots, the right-half, split violin plots show the probability density, and vertical black lines show the median, IQR, hinges showing the first and third quartiles, and vertical whiskers showing +/- 1.5IQR. The binned dotplots on the left half show each individual subject as a point. **B. Effect of condition on accuracy**. We found a significant effect of the condition (which stimulus was stronger) on accuracy, despite matched available posterior information, suggesting participants to give suboptimal relative weights to priors and likelihoods in their decisions. Participants performed significantly better in the Stronger-Target condition (M_L_Stronger-Target_=0.72, SD=0.05; M_M_Stronger-Target_ =0.79, SD=0.04; M_H_ Stronger-Target_ =0.84, SD=0.04) compared to the Stronger-Lead condition (M_L_Stronger-Lead_=0.70, SD=0.05; M_M_Stronger-Lead_=0.76, SD=0.05; M_H_Stronger-Lead_=0.81, SD=0.05), pointing to a relative underweighting of prior information. **C. Effect of condition on confidence**. We found no effect of condition on confidence following correct trials (with mean differences between conditions – Stronger-Target minus Stronger-Lead – of: M_L_=0.15, SD=2.19; M_M_=0.40, SD=2.34; M_H_=-0.07, SD=2.15). But, we found an effect of condition following incorrect trials, with lower confidence in the Stronger-Lead condition (with mean differences between conditions – Stronger-Target minus Stronger-Lead – of: M_L_=0.66, SD=2.51; M_M_=1.98, SD=2.50; M_H_=3.15, SD=2.54). From the decision level as a baseline, this result indicates greater use of the prior in confidence than in decisions. **D. M-Ratio estimates**. M-Ratios per participant for each condition, measured across all posterior levels. As expected if (as panels B. and C. suggest) valid prior information informed confidence ratings more than discrimination decisions, M-Ratios were significantly higher in the Stronger-Lead condition than in the Stronger-Target condition.

#### Dependence on Condition Reveals Suboptimal Prior Weighting in Decisions

After confirming that the manipulations worked as intended, we investigated the use of prior information in decisions. We reasoned that, if the prior were suboptimally weighted at the decision level, performance on the target decision would depend on the condition despite matched posterior information (Figure 1C). To evaluate this, we built a logistic mixed effects model on target decision response accuracy. This included fixed effects of posterior level (L, M, H), condition (Stronger-Lead, Stronger-Target), and their interaction (Table 1). We did not find a significant interaction effect. In line with our manipulation check, we found a significant main effect of posterior level (*χ*^2^(2)=175.77, p<0.001, BF_10_=6.12 × 10^36^), with accuracy increasing with higher available posterior information (Figure 2B). We also found a significant main effect of condition, *χ*^2^(1)=9.06, p=0.003, BF_10_=8.95, with better performance when the target was stronger, OR_Stronger-Lead/Stronger-Target_=0.89, 95% CI [0.82, 0.96], Z=-3.09, p=0.002 (Figure 2B). This suggests that, despite an equal amount of available posterior information, participants performed better when more of that information was carried by the target stimulus, rather than the lead stimulus, indicating that they dismissed some of the prior information when making their decisions. In other words, this revealed an underweighted use of the prior at the decision level, relative to an optimal observer. Because there was potential for response bias to lead to differences in accuracy across the conditions regardless of how the prior information was weighted, we reran this model, including a measure of response bias obtained from fitting subject-wise psychometric functions to performance on the unbiased first decisions (see Methods). This did not substantially change the result or conclusions, as there were no significant effects involving response bias.

#### Analysis of Mean Confidence Reveals Greater Use of Priors Than in Decisions

We then built a linear mixed effects model on confidence with fixed effects of posterior level (L, M, H), condition (Stronger-Lead, Stronger-Target), target response accuracy (Correct, Incorrect), and their interactions (Table 1). We found evidence for a three-way interaction from Bayesian statistics BF_10_=70.83 (Figure 2C) but only a trend from frequentist tests (F(2,14832)=2.66, p=0.070). Post-hoc pairwise comparisons revealed a significant effect of condition on confidence only for incorrect trials at the high posterior level, t(1281)=-3.52, p<0.001, BF_10_=6.14, η^2^_p_=0.01, and medium posterior level, t(839)=-2.16, p=0.03, η^2^_p_=0.006, although the Bayesian statistics suggest evidence against the effect at the medium level (BF_10_=0.46). There were no significant effects of the condition following correct trials, with evidence for the null hypothesis at all posterior levels (all BF<0.02). Because the suboptimal weighting of the prior at the decision level can impact confidence distributions, a statistically significant effect of condition on confidence cannot be interpreted in isolation. Instead, we must consider the decision level result as a baseline. Simulations shown below (Figure. 4A, third panel) revealed that, given any underweighting of the prior in decisions, the following pattern holds: If the prior was even further underweighted in confidence than in decisions, we would expect confidence in the Stronger-Target condition to be higher following correct trials and lower following incorrect trials. Contrary to this, the result of no difference following correct trials and lower confidence in the Stronger-Lead condition following incorrect trials indicates that the prior is less underweighted than at the decision level, and is hence used more optimally in confidence. To quantify this effect and concretely compare the weighting of the prior at the decision versus confidence level, we used computational modelling. Further in line with this idea, we found significantly higher M-Ratios in the Stronger-Lead condition (M=0.87, SD=0.27) than in the Stronger-Target condition (M=0.71, SD=0.25), t(16)=3.64, p=0.002, η^2^_p_=0.01, 95% CI [0.14, 1.00] (Figure 2D), suggesting that the relative use of priors and likelihoods is more optimal at the metacognitive level than at the level of decisions.

#### Bayesian Model Fits Different Weights of Priors in Confidence and Decisions

Overall, the behavioural analyses revealed that, while decisions underweighted the available prior information, confidence seemed to use the prior information more optimally. To account for these results, and to be able to compare the effects at these two levels of processing more directly and quantitatively, we use a computational model. We built and fit a Bayesian model of decisions and confidence under informative priors that included a weighting parameter for the prior (relative to the likelihood), for both the decision (*w*_*choice*_) and the confidence rating (*w*_*conf*_). These weighting parameters reflected potential over- or underestimation of the precision of the prior, hence weighting its influence over behaviour relative to the likelihood. They were also computationally identical in their impact, scaling the perceived variance of the prior, allowing us to directly compare them. We first estimated internal noise and decision bias (not under the influence of a prior) per participant on separate data using the Akaike-weighted combination of four fit psychometric functions (see Methods), and then incorporated these subject-wise estimates into the model, allowing us to account for them. The model additionally included a parameter for confidence bias (*b*) that captured overall over- or underconfidence in both stimuli. The model had three free parameters (per participant and at the group level): *w*_*choice*_, *w*_*conf*_, and *b*. We confirmed the parameters to be recoverable in a parameter recovery analysis (Supplementary Materials, Figure S2B). We also compared our full model, which we call the Flexible model, to a variety of simpler models, to assess each main research question in more depth.

#### Flexible Model Results

At the decision level, the model predicted that if the prior was optimally weighted relative to the likelihood, (*w*_*choice*_ = 1), there would be no difference in accuracy between the two conditions (Figure 3A, left panel). If the prior was overweighted (*w*_*choice*_ < 1, capturing an underestimation of the variance of the prior), accuracy would be higher when the lead was the stronger stimulus (Figure 3A, middle panel). Conversely, if the prior was underweighted (*w*_*choice*_ > 1), accuracy would be higher when the target was the stronger stimulus (Figure 3A, right panel). The full hierarchical model fit the data with a mean of the posterior for the group mean parameter *w*_*choice*_ = 2.17 (SE=0.007; Figure 3C). This suggests, in line with the behavioural results, that priors are underweighted in decisions, relative to an optimal observer. The simulated choice behaviour for these model results is shown in Figure 3B (right panel) and demonstrates higher accuracy when the target is the stronger stimulus, just as we found behaviourally.

**Figure 3.**
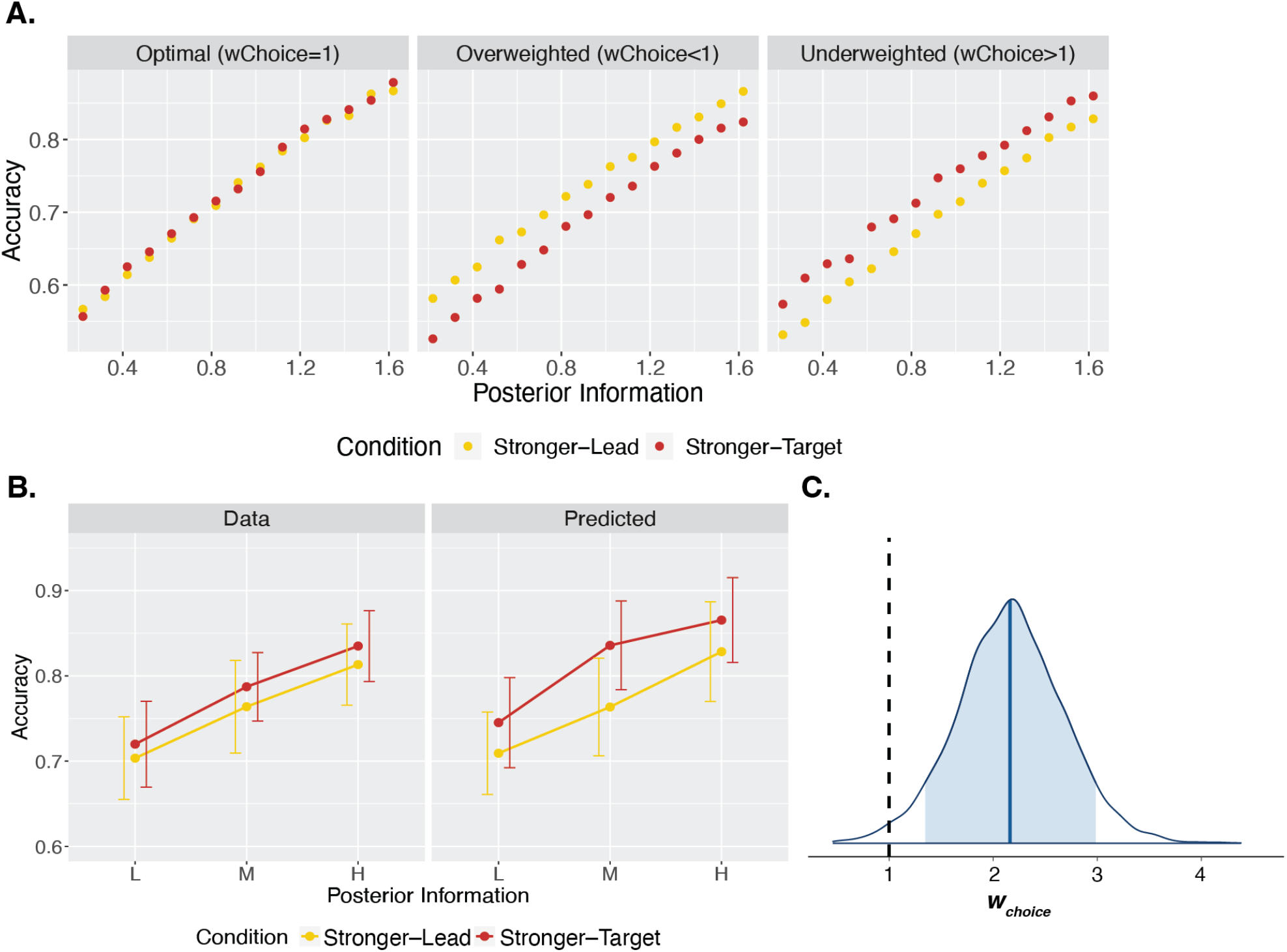
Modelling Discrimination Decisions. **A. Model simulations of decision accuracy**. Target decisions were simulated from the Flexible model across different posterior levels, between the two conditions, and at three different values of *w*_*choice*_: 1 (optimal weighting of prior information), 0.33 (overweighting of prior), and 3 (underweighting of prior), shown from left to right respectively. These values are representative of the range found in the data, and capture either over- or underestimated variance by a factor of 3. The resulting decision accuracies are shown here. The model predicts that optimally using prior information will lead to no difference in accuracy between the two conditions, whereas overweighting prior information will lead to higher accuracy when the lead is stronger, and underweighting prior information will lead to higher accuracy when the target is stronger. **B. Data and predictions of the fit model**. The left panel shows the observed accuracies per posterior level and condition. The right panel shows the predicted accuracies generated from sampling the fit posterior group mean parameter distributions 1000 times, and simulating 720 trials per participant for each of those sampled parameters. Note that we use the sample group mean parameter for simulating trials, but still used each participant’s staircased coherences, internal noise and decision bias. Error bars capture standard deviation (SD) of accuracies across participants. **C. Posterior distribution for *w***_***choice***_. The posterior distribution for the group mean parameter of the weighting of prior information in the decision, *w*_*choice*_. The blue shaded region shows the 89% credible interval and the vertical black dashed line reflects optimal weighting of the prior in the decision (*w*_*choice*_ = 1). A weight above 1 captures overestimation of the variance and hence underweighting of the prior.

We then examined the model predictions for confidence, given the result of the underweighted priors in decisions (*w*_*choice*_>1). With overweighting of the prior in confidence (*w*_*conf*_<1), the Stronger-Lead condition tends to show higher confidence than the Stronger-Target condition for correct trials, and lower for incorrect trials (Figure 4A, middle-left panel). In other words, the differences in mean confidence between correct and incorrect are larger in the Stronger-Lead condition than in the Stonger-Target condition. The model predicts these differences between conditions to become more extreme as the prior is weighted more strongly in confidence (as *w*_*conf*_ decreases). In contrast, with underweighting of the prior (*w*_*conf*_>1), it is the Stronger-Target condition that tends to have larger differences in mean confidence between correct and incorrect (Figure 4A, middle-right panel). Again, the model predicts this pattern to get more extreme as the prior is weighted more weakly (as *w*_*conf*_ increases). If the prior weighting in both decisions and confidence were optimal, the model predicts no difference in mean confidence between conditions. However, due to the suboptimal weighting found at the decision level, the model predicts differences between conditions with optimal *w*_*conf*_ = 1, shown in Figure 4A, far-left panel. The behavioural pattern seen in confidence — of no difference between conditions following correct trials, and lower confidence following incorrect trials in the Stronger-Lead condition — can only be produced if the prior is weighted more strongly in confidence than in the decision, and the model predicts such a pattern when the prior is underweighted, but less so than in decisions (Figure 4A, far-right panel).

**Figure 4.**
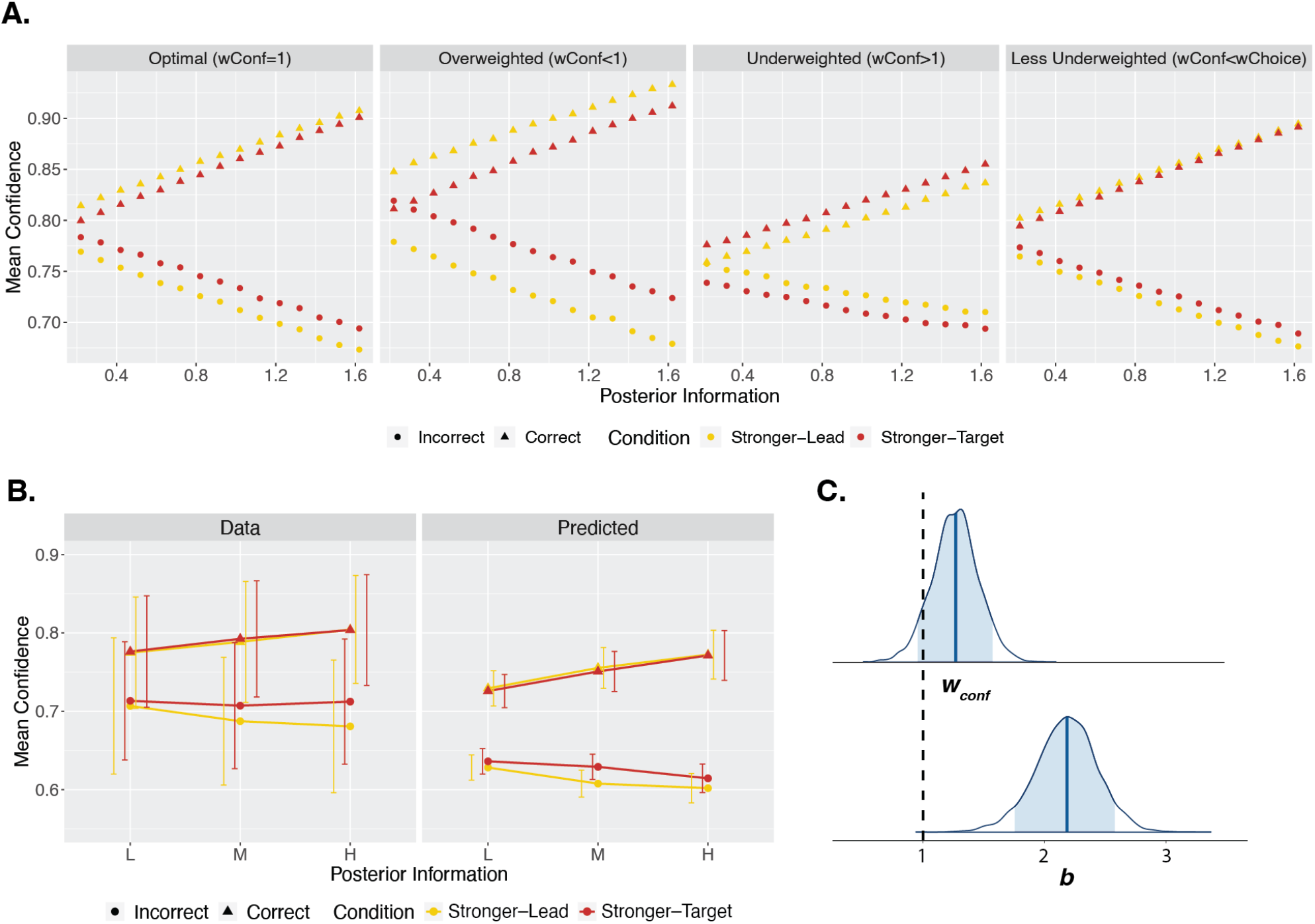
Modelling Confidence. **A. Model simulations of confidence**. Confidence ratings following correct and incorrect trials were simulated from the model across different posterior information levels, between the two conditions. Simulations used the *w*_*choice*_ value from the model fit, 2.17, and four different values of *w*_*conf*_: 1 (optimal weighting of prior), 0.33 (3-fold overweighting of the prior), 3 (3-fold underweighting of the prior), and 1.27 (the value obtained from the model fit, underweighting the prior in confidence less than in the decision), shown from left to right respectively. The resulting mean confidence values are shown here. The model predicts mean confidence to increase with increased available posterior information following correct decisions and decrease with increased available posterior information following incorrect trials. As the prior is increasingly overweighted, the model predicts higher mean confidence in the Stronger-Lead condition following correct decisions, and lower in the Stronger-Lead condition following incorrect decisions. As the prior is increasingly underweighted, the opposite pattern is predicted, with higher mean confidence in the Stronger-Target condition following correct trials, and lower confidence in the Stronger-Target condition following incorrect trials. If the prior weighting was optimal in both decisions and confidence, there would be no differences predicted in mean confidence between conditions. However, due to the suboptimal weighting at the decision level, the model predicts differences in mean confidence when *w*_*conf*_ is optimal, as shown in the far-left panel. When the prior is underweighted in confidence, but less so than in the decision, the model can produce the pattern seen behaviourally, in which there is no difference between conditions following correct trials, but lower confidence in the Stronger-Lead condition following incorrect trials. **B. Model results against data**. On the left are the data, showing confidence following correct and incorrect decisions per posterior level and condition. On the right is mean confidence generated from sampling the fit posterior parameter distributions 1000 times and simulating 720 complete trials per participant based on those parameters. Note that we use the sample group mean parameters for simulating trials, but still used each participant’s staircased coherences, internal noise and decision bias. The error bars capture standard deviation (SD) of the mean confidence across participants. **C. Posterior distribution for *w***_***conf***_ **and *b***. The top posterior distribution is for the group mean parameter of *w*_*conf*_. The lower posterior distribution is for the group mean parameter of *b*. The blue shaded regions show the 89% credible intervals and the vertical black dashed line corresponds to the parameter values of an optimal observer.

Due to complexity constraints, we modelled confidence on each trial assuming that the internal signal corresponded to the mean external stimulus strengths from each stimulus coherence level (L, M, H). That is, we did not fit the internal latent samples that are theoretically generated from the external stimuli, as that would have led to an overparameterized model with two free parameters per individual trial (totalling close to 30000 free parameters). This simplification still included a parameter that corresponded to the internal noise for each participant, which was used in the confidence computation (see Methods). As a result of this simplification, the model underestimated mean confidence and impacted primarily the confidence bias parameter *b*. However, this still allowed the model to capture differences between conditions, and hence our parameter of interest, *w*_*conf*_. Additionally, we confirmed that any impact that this had on *w*_*conf*_ worked directly against our conclusions (see Supplementary Materials), making our interpretations more conservative.

The fit hierarchical model had a mean posterior of the group parameter *w*_*conf*_ = 1.27 (SE=0.002), and of the group bias *b* = 2.18 (SE=0.003; Figure 4C). This, in line with the behavioural results, suggests the weighting of the prior in confidence to be closer to optimal, compared to the weighting of the prior in decisions. The simulated confidence for these model results at the group level (Figure 4B) successfully captures the pattern associated with the combination of an underweighted prior in decisions and a less underweighted prior in confidence (Figure 4A, far-right panel).

Finally, we directly compared the relative weighting of priors and likelihoods at the decision and confidence level. *w*_*choice*_ was credibly different from *w*_*conf*_, with the difference distribution just excluding 0 in the 89% credible interval [0.03, 1.77] (Figure 5B). This again suggests the weighting of priors relative to likelihoods to differ at the level of decisions and confidence.

**Figure 5.**
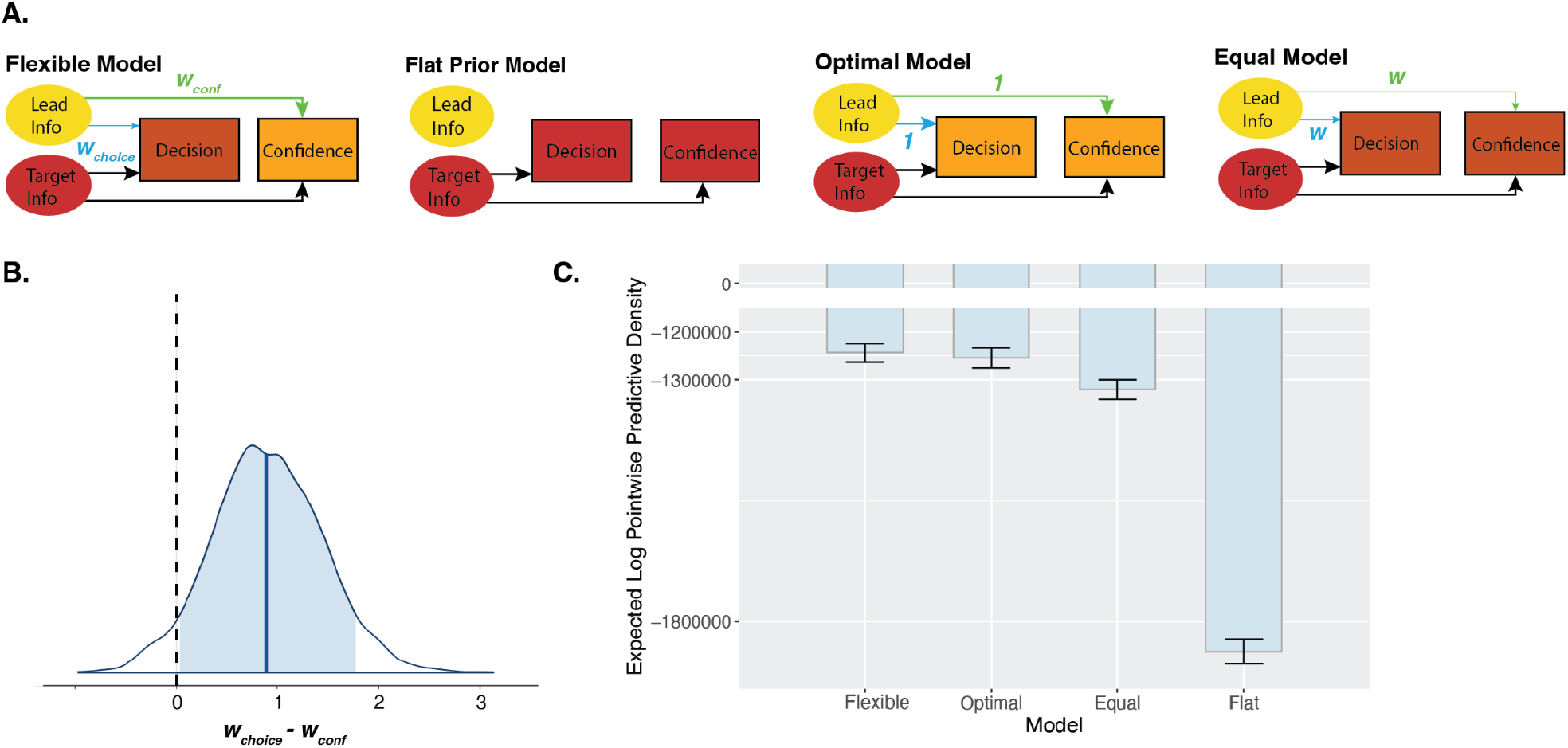
Model Comparison Results. **A. Schematic of the Four Compared Models**. Schematics of the four models we compared, with the thickness of the light blue arrows depicting the weighting of priors relative to likelihoods in the decision and the thickness of the light green arrows depicting the weighting of the priors relative to likelihoods in confidence. In the Flexible Model, relative weighting of priors and likelihoods was allowed to differ between choices and confidence (w_choice_ was allowed to differ from *w*_*conf*_). The Flat Prior Model only has arrows from the target, showing that the prior information was not used at all in decisions or confidence. In the Optimal Model, *w*_*choice*_ and *w*_*conf*_ both had to be optimal and equal to 1. In the Equal Model, the weighting of priors relative to likelihoods was allowed to differ from optimal but the pattern was the same between *w*_*choice*_ and *w*_*conf*_. **B. Posterior Group Difference Distribution of *w***_***choice***_ ***-w***_***conf***_. The posterior distribution for the difference in the group mean parameters *w*_*choice*_ and *w*_*conf*_. The blue shaded region shows the 89% credible interval and the vertical black dashed line reflects no difference in the two parameters (*w*_*choice*_ - *w*_*conf*_ = 0). 0 is just excluded from the 89% credible interval, suggesting *w*_*choice*_ and *w*_*conf*_ to be credibly different from one another. **C. Expected Log Pointwise Predictive Density (ELPD) Results from LOGO-CV**. The predictive capacity of each model is shown as the elpd value from the LOGO-CV with 10 folds, leaving out and then predicting 2 participants per fold. This shows the Flexible Model to have the highest predictive capacity, suggesting it as the best model to explain the data. This is followed by the Optimal Model, Equal Model and then Flat Prior Model. Error bars depict SE.

#### Flexible Model Describes Behaviour Better Than Simpler Models

We then compared the Flexible model to simpler models (Figure 5A). We confirmed these models to be distinguishable from one another with a model recovery analysis (see Supplementary Materials, Figure S4).

In addition to the (1) Flexible model, we built (2) the Flat Prior model in which the lead stimulus information was not used at all and the target stimuli were considered to occur under a flat prior (*w*_*choice*_ and *w*_*conf*_ infinitely large), (3) the Optimal model in which *w*_*choice*_ and *w*_*conf*_ were both optimal (equal to 1), and (4) the Equal model in which the prior was forced to be used to the same extent in decisions and confidence (*w*_*choice*_=*w*_*conf*_) and hence only one *w* parameter was fit. All models included confidence bias as a free parameter. We fit these models and compared their predictive performance against the Flexible model using a 10-fold leave-one-group-out cross-validation (LOGO-CV, where ‘groups’ correspond to participants). The Flexible model predicted the data better than each of the other models, with a difference in expected log pointwise predictive density (elpd_diff) of -619688.2 (se_diff=16717.0) compared to the Flat Prior model, of -11017.7 (se_diff=4534.4) compared to the Optimal model, and of -76685.8 (se_diff=2147.4) compared to the Equal model (Figure 5C). The results of this formal model comparison are in line with both the behavioural and the modelling results shown above: The difference to the Flat Prior model confirmed that participants used the prior information to some extent, in line with our behavioural manipulation check. The difference to the Optimal model confirmed that participants used the prior information suboptimally. And, the difference to the Equal model confirmed that participants used the prior information to a different extent in confidence than in decisions, in line with our finding of a credible difference between *w*_*choice*_ and *w*_*conf*_.

### Experiment 2

#### Dual-Decision Task with Delayed Target Decision

The finding from Experiment 1, that priors can be used more optimally in confidence, might support the idea that priors are integrated gradually, and that there is continued post-decisional evidence accumulation that can then factor into confidence. This would be in line with previous work suggesting that confidence computations incorporate additional information that has accumulated after the first-order decision^20–23,25,29,30^. Alternatively, it is possible that the additional use of the prior in confidence is by virtue of the introspective act, and not simply due to continued evidence accumulation. In support for the latter alternative, previous work found enhanced metacognitive efficiency following prior congruency, showing information from priors to especially boost metacognitive judgments^27^. In order to investigate these two possibilities further, we ran a second pre-registered experiment (N=25) in which we repeated the same paradigm under the same conditions, but we added a 2-second delay after the offset of the target stimulus and before participants were allowed to report their target decision. We chose the duration of the delay to approximately match the peak of the distribution of reaction times between viewing the target stimulus and giving the confidence rating from Experiment 1, which was 2.48 seconds. Therefore, if in Experiment 1 the more optimal use of the priors in confidence was only due to the extra processing time before giving the confidence rating, then delaying the target decision until that time point in Experiment 2 should lead to more optimal use of the prior information in the delayed target decision. If, however, the more optimal use of the prior was due to the introspective confidence rating, delaying the target decision in Experiment 2 should not change the pattern of results.

#### Priors Underweighted in Decisions Despite Increased Processing Time

We ran the same regression model on target response accuracy as in Experiment 1 and found a significant interaction between posterior level and condition, *χ*^2^(2)=8.21, p=0.017, BF_10_=2.68. In line with Experiment 1, response accuracy was higher in the Stronger-Target condition at each posterior level, although this was only significant at the medium posterior level, z=-4.30, p<0.001, BF_10_=138.49 (Figure 6A). This suggests the prior to be underweighted in the target decisions, even after the delay and hence the added opportunity for evidence accumulation (for behavioural data, see Supplementary Materials, Figure S5). To examine whether the same pattern of more optimal use of priors in confidence remained, we again fit the Flexible model, this time as confirmatory modelling analyses. Replicating the pattern found in Experiment 1, we found a mean of the posterior of *w*_*choice*_ = 3.97 (SE=0.02; Figure 6A) and a mean of the posterior of *w*_*conf*_ = 1.39 (SE=0.01; Figure 6B), with a credible difference between them [0.50, 4.64] (Figure 6C). Taken together, this suggests that prior information is underweighted in decisions even when those decisions occur at a similar time point as the confidence judgments in Experiment 1, and, like in Experiment 1, confidence then more optimally uses that prior information.

**Figure 6.**
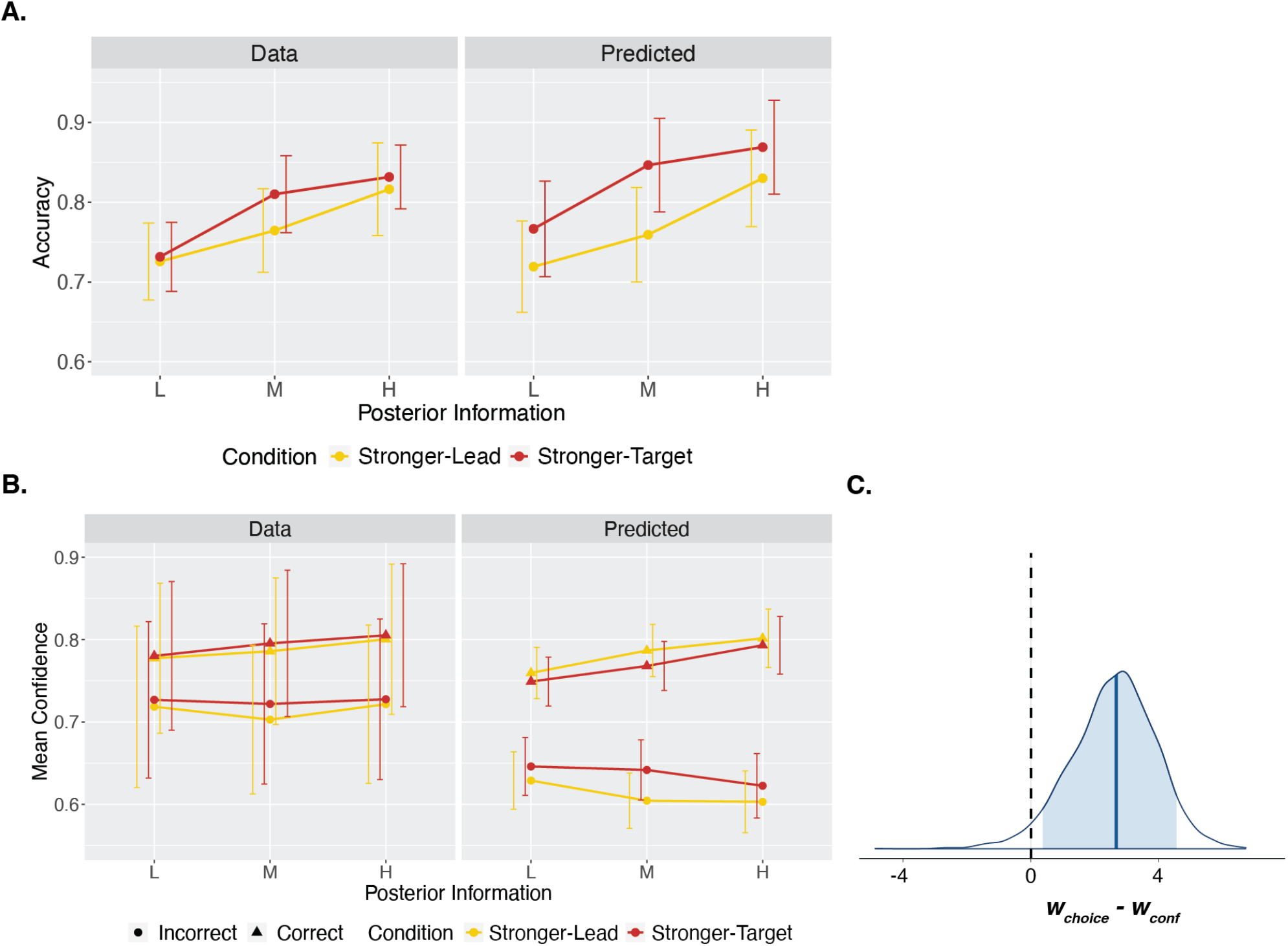
Experiment 2 Results. **A. Decision level model results against data**. The left panel shows the observed accuracies per posterior level and condition. The right panel shows the predicted accuracies generated from sampling the fit posterior group mean parameter distributions 1000 times, and simulating 720 trials per participant for each of those sampled parameters, also using each participant’s staircased coherences, internal noise and decision bias. Error bars capture standard deviation (SD) of accuracies across participants. **B. Confidence level model results against data**. On the left are the data, showing confidence following correct and incorrect decisions per posterior level and condition. On the right is mean confidence generated from sampling the fit posterior parameter distributions 1000 times and simulating 720 complete trials per participant based on those parameters, also using each participant’s staircased coherences, internal noise and decision bias. Error bars capture standard deviation (SD) of mean confidence across participants. **C. Posterior Group Difference Distribution of *w***_***choice***_ ***-w***_***conf***_. The posterior distribution for the difference in the group mean parameters *w*_*choice*_ and *w*_*conf*_. The blue shaded region shows the 89% credible interval and the vertical black dashed line reflects no difference in the two parameters (*w*_*choice*_ - *w*_*conf*_ = 0). 0 is excluded from the 89% credible interval, suggesting *w*_*choice*_ and *w*_*conf*_ to be credibly different from one another.

## Discussion

In two experiments, we tested whether prior information influences confidence optimally, and how this compares to its influence on perceptual decisions. To do so, we compared pairs of conditions that were matched in the available posterior information but differed on whether the stronger source of information was the prior or the new incoming information. We then evaluated the differences between conditions (Stronger-Lead vs Stronger-Target) in both discrimination accuracy and mean confidence, and fit a quantitative model to measure the weighting of prior information. This revealed that priors are underweighted relative to likelihoods in discrimination decisions. Conversely, confidence judgments incorporated prior information to a greater extent than discrimination decisions did. Further, and in line with the idea that prior information is processed more optimally at the metacognitive level, we found that metacognitive efficiency was higher when more information was carried by the prior, with first-order performance hindered while confidence preserved the use of this information. Taken together, these results suggest that we can access and use information from priors in explicit, introspective confidence judgments even to a greater extent than we use that information to guide decisions.

This pattern goes against the assumptions, implicit in the Bayesian framework, of optimal and equal integration of priors in decisions and confidence. While participants may not necessarily be expected to behave as Bayesian optimal observers, these findings quantify precisely in which way they deviate from those assumptions. Although the underweighted prior in decisions may, in isolation, be explainable by a decay of the prior information over time, such a decay would make the asymmetry between the decision and confidence levels even more surprising, as the confidence judgments occurred even later. The results of Experiment 2 revealed, further, that this asymmetry remains even when additional processing time is given by forcing a delay before the target decision. This suggests that this pattern cannot be accounted for just by continued evidence accumulation before the confidence rating, since a similar amount of evidence accumulation should have occurred between the confidence ratings in Experiment 1 and the target decisions in Experiment 2. Instead, this points towards a more optimal use of prior information at the metacognitive level, compared to first-order processing. In line with this, Balsdon and colleagues demonstrated asymmetries in the information used by decisions and confidence^24,25^. They used series of stimuli and found decisions to set covert bounds and stop collecting new evidence, while confidence used more of the available information. In light of those findings, our work shows a similar effect, where decisions make use of less of the information available than confidence does. However, here, we find that it is prior information that is more strongly dismissed in decisions.

These results add a novel layer to recent findings by Lisi et al.^28^, who focussed on implicit, not explicit confidence. They, as we did, found priors to be underweighted at the level of the decision, but could not assess whether explicit confidence weighted them differently than decisions. Here we reveal important differences in how the prior is used at different processing levels, by examining the weighting of the prior in explicit confidence as well. Our results suggest that, even though the prior is underweighted in a decision, people can access and use this information better when asked to make an explicit introspective judgment about that decision. A cognitive architecture in which perceptual decisions can primarily respond to current incoming evidence while higher order metacognitive processing integrates different sources of information and monitors their relative certainty might be highly adaptive. For example, it might be beneficial to react rapidly and in accordance with evidence for even an unlikely belief if that would pose some threat, meanwhile having the metacognitive system accurately track its posterior probability for appropriate models of the world. Our results therefore provide crucial insight on a dissociation between human behaviour and associated confidence.

In this study, we used a paradigm with informative, high level priors. Future work is necessary to investigate whether this result holds true when different kinds of priors are used. First, lower-level priors such as the light-from-above prior, cardinal orientation bias, or perceptual history bias might affect decisions differently, as they may act at an earlier stage and impact perception of the target stimulus more directly^31–33^. Second, non-informative or suboptimal priors might reveal that the pattern we see here reflects a confidence bias towards prior information, rather than more optimal use of priors in confidence. If so, in cases of suboptimal priors, confidence would still be more likely to be affected by the invalid prior information than decisions. This possibility is in line with recent studies that have shown that confidence is biased by suboptimal, false priors about stimulus precision^34^ or about task performance^35^. Other work testing the Bayesian confidence model has found confidence to suboptimally overweight evidence that is in line with the decision, leading to a form of confirmation bias in perceptual confidence^17,36,37^. Our findings do not show a confirmation bias in favour of information in line with the decision, but might rather reflect a confidence confirmation bias in favour of information in line with the prior, even in cases where this actually contradicts the decision. Although at face value this may go against the previous confirmation bias findings in perceptual confidence, this might, speculatively, still be in line with the conclusions drawn, namely that confidence favours evidence consistent with one’s beliefs. This could also be a strategy aimed at self-consistency and avoiding cognitive dissonance^37–39^, leading people to be more confident in response to information that fits in their belief system, and to doubt themselves when they act against their prior world models.

## Methods

Both experiments were pre-registered (Experiment 1: https://osf.io/qgpsr and Experiment 2: https://osf.io/tvyrz), and we respected the pre-registered plan unless stated otherwise.

### Participants

For Experiment 1, we pre-registered that we would test 25 participants across two sessions. We chose this sample size to be close to previous studies using similar tasks and modelling methods^8,28^, which included between 15 and 26 participants. We also pre-registered six minimal criteria to invite participants to the second session. The most important of these criteria were (1) that response accuracy increased across the three coherence levels — hence suggesting that the experimental manipulation had the intended effect on internal signals —, and (2) that response accuracy was (any amount) higher on the second decisions as compared to the first, indicating basic use of the task structure. Following these criteria, we excluded 12 participants without inviting them to take part in the second session, and collected data until we reached 25 participants that met these criteria and were tested for two sessions. Four of these participants were later excluded from analysis because they no longer met these basic criteria after including data from their second session, leaving a total of 21 participants (10 male, 11 female) included in the analyses. Participants were tested in Berlin, were healthy and were between 18 and 37 years of age (M=25.7, SD=4.6). Participants all reported to have normal or corrected-to-normal vision, were fluent in English, and primarily right handed (Edinburgh Handedness Inventory score: M=83.2, SD=28.5). Participants were compensated with 8€ per hour or with equivalent course credit and gave signed, informed consent before starting the experiment. The ethics committee of the Institute of Psychology at the Humboldt-Universität zu Berlin approved the study (Nr. 2021-47), which conformed to the Declaration of Helsinki.

For Experiment 2, we pre-registered that we would test an initial 25 participants that met the minimal exclusion criteria after both sessions, after which we set a stopping rule, based on evidence for or against the effect of condition on target response accuracy. After 25 participants, we found substantial evidence for the alternative hypothesis and stopped collecting data. These 25 participants included 9 male, 15 female, and 1 that did not specify; were between 19 and 34 years of age (M=25.4, SD=3.8); and were primarily right handed (Edinburgh Handedness Inventory score: M=86.3, SD=34.3 - note one participant was excluded from this due to missing data), as well as meeting the same inclusion criteria as in Experiment 1.

### Setup

The experiment was programmed in HTML/Javascript/CSS to run in the browser. We used JATOS^47^ to store the result data. The study ran on Google Chrome (version 94.0.4606.71) on a Dell Precision 5760 laptop (Intel core i7 with 31GB of RAM) with a display resolution of 1,920 × 1,200 (refresh rate=60Hz).

### Procedure

#### Control Task

In each of the two sessions, prior to starting the main task, participants first completed 90 trials of a control task. Each trial of the control task consisted of a single dot motion stimulus with a 50% chance of the coherent motion going to the right vs left, followed by a right/left decision, which participants made using the “S” or “A” keys, respectively. The stimuli in the control task spanned six different coherences, meant to capture a broad range of difficulties - 5%, 10%, 12%, 15%, 20% and 30% coherence. The resulting data were later used to estimate participants’ internal noise and decision bias (see below). In total across the two sessions, participants completed 180 control task trials (30 per coherence level). After the control task, we explained the instructions for the full task structure to participants both verbally and in written instructions, and they then completed five demo trials to familiarise themselves with the task and buttons, and then proceeded to the main task.

#### Main Task

On each trial of the main task, participants completed two consecutive decisions consisting of a random dot motion stimulus followed by a right/left decision using the “S” or “A” keys (Figure 1A). The first (lead) stimulus of each trial had a 50% chance of the coherent motion going to the right vs left. The direction of coherent motion of the second (target) stimulus depended deterministically on the response accuracy of the first decision such that if they were correct, the second stimulus would have coherent motion to the right, and if they were incorrect, it would have coherent motion to the left. Participants were informed of this rule and instructed to use this to help them in the task. Following an optimal strategy, this conditional rule meant that participants should expect a rightward second stimulus with a strength of prior expectation proportional to their decision confidence about the first decision. Following the second decision, participants rated their confidence on a continuous sliding scale from 50% (guessing) to 100% (totally sure), using the mouse. The dual-decision task structure with the conditional rule was an extension of a previous study investigating implicit confidence^28^.

The task was gamified to make it more engaging. The background of the screen was an illustration of fields with a barn to the right, and the stimulus display circle in the middle of the screen (Figure 1A). The first decision controlled the movement of a cartoon sheepdog to the right or left of the stimulus display, and the second decision controlled the movement of a cartoon farmer to the right or left of the stimulus display. We explained to participants that the moving dot stimuli depicted flocks of sheep, with some “leader” sheep that moved coherently to either the left or right, and that they had to decide based on the motion direction of the leader sheep whether to send their sheepdog to the left or right. Participants were explicitly informed of the rule: If they were correct, and hence the sheepdog was in the correct place, the sheep would be herded toward the barn and on the next stimulus, they would be going to the right. If, however, they were incorrect and the sheepdog was not in the correct place, the sheep would run away and on the next stimulus, they would be going to the left. After the second stimulus, participants then had to make the final decision to either send the farmer to the barn (to the right) to get the sheep, or to send the farmer to herd them from the fields to the left. They then rated their confidence that the farmer had successfully gotten the sheep. We emphasised to participants that they should use the rule to try to help them with the task, and that they should take time to give as sensitive and meaningful confidence ratings as possible.

We manipulated the coherences of the stimuli to create three stimulus levels, and the coherence levels of each of the two stimuli per trial combined to form three overall posterior levels. The low posterior information level consisted of one low and one medium coherence stimulus (L+M or M+L), the medium posterior information level consisted of one low and one high coherence stimulus (L+H or H+L), and the high posterior information level consisted of one medium and one high coherence stimulus (M+H or H+M). These posterior information levels existed across two conditions, a “Stronger-Lead” condition in which the lead stimulus was stronger, and a “Stronger-Target” condition in which the target stimulus was stronger. The stimulus coherence levels were staircased by staircasing lead stimuli, which were not under the influence of an informative prior, with the medium level staircased via a 2-down-1-up procedure targeting 71% accuracy, and the high level staircased via a 3-down-1-up procedure targeting 79% accuracy. The low level was yoked to the medium staircase, but remained 5% lower in coherence, as there was no N-down-1-up procedure that would target an accuracy between 50% and 71%. The three posterior information levels as well as the condition were counterbalanced across each block. Participants received feedback about their performance on the target decisions at the end of each block. Each block consisted of 36 trials, and participants completed 10 blocks per session for a total of 360 trials per session and 720 trials in the experiment. Each session took between 1-1.5 hours in total.

#### Experiment 2

The paradigm remained the same in Experiment 2 except that there was an added delay period of 2 seconds before participants could enter the target decision using the “S” or “A” key, after viewing the target stimulus. After these 2 seconds, a light grey ring appeared around the viewing circle to indicate to participants that they could now report their decision. Participants were instructed to try and avoid pressing a key prematurely during the delay period, although trials with premature presses were not excluded. Participants received feedback at the end of each block about how many premature presses were made, in order to remind them to limit this.

#### Stimuli

The dot motion stimuli were made using an adapted version of an RDK jsPsych plugin^48^. Stimuli were composed of 100 total moving white dots on a circular gray background. Each dot had a radius of 2 pixels and the background circle had a diameter of 425 pixels, with an aperture diameter of 319 pixels (75% of the circle diameter). The noise dots had constant directions that were randomly sampled, and the coherent dots moved in a constant horizontal direction either to the left or right. All dots moved 2 pixels per frame and had a dot life of 17 frames (i.e., each dot followed their trajectory for 17 frames before being redrawn at a random location). Each stimulus was presented for 300 ms. Although some directional information was possible in the random dots of each stimulus, we confirmed that this did not lead to an overall bias in any participant, so that stimulus directions remained balanced between left and right (for the lead decision, where they were intended to be 50/50), even with the directional information from the noise dots. None of the decisions or confidence ratings was speeded.

### Analysis

We removed any trials with reaction times longer than 8 seconds on any decision or confidence ratings.

Our main behavioural hypotheses were tested using the ‘lme4’ package^49^ in R^50^ for building linear and generalised linear mixed-effects models. For each regression analysis, we used the most complex random effects structure that converged on the full model^51^, which meant deviating from pre-registered random effects for analysing confidence. Model syntaxes can be seen in Table 1. All hypotheses were tested using two-tailed tests and an alpha level of 0.05, and reported *χ*^2^ values are based on a comparison of the model of interest and null model with the same random-effects structure. We computed effect sizes for the linear mixed-effects analyses as η^2^_p_ and reported 95% confidence intervals whenever they were available. We additionally computed Bayes factors for our main hypotheses using the ‘BayesTestR’ package^52^ and using Bayesian models with uniform priors with the ‘brms’ package^53^. For these Bayesian regressions, we ran 4 chains of 10000 iterations, including 2000 burn-in samples, which gave a total of 32,000 effective samples, and we ensured a R-hat close to 1. To analyse whether the prior was suboptimally weighted at the decision level, we deviated from our pre-registered regression approach of examining the effect of condition on the probability of choosing “right” given rightward stimuli. We realised from later simulations that this would not sufficiently distinguish between an optimal and suboptimal weighting of the prior. We instead examined the effect of condition on response accuracy, which could adequately address this question.

In order to compare participants’ metacognitive efficiency between conditions, we used the M-Ratio measure (meta-d’/d’) described in previous work^54^, with R scripts available from https://github.com/craddm/metaSDT. Two participants were removed from this analysis due to extreme confidence distributions, with over 40% of trials at 100% confidence. Two further participants were removed due to Type 1 hit rates above 0.95 in either condition, but the results did not change when these two participants were included. For measuring M-Ratio, we transformed participants’ continuous confidence ratings to a 5-bin discrete scale using quantiles, computed on all ratings per participant.

### Modelling

To quantitatively assess how participants weighted the prior in their decisions and confidence, we fit a Bayesian model to their data. The specific model definitions, model fitting, model selection and evaluation of model results were all exploratory and the details were not pre-registered. The full model, which we refer to as the Flexible model, included two free weighting parameters, *w*_*choice*_ and *w*_*conf*_, that captured the weighting of the prior relative to the likelihood in the decision and in confidence, respectively. These parameters acted in the same way in the model, scaling the estimate of the variance of the prior, and could hence be directly compared. The model also took as input a measure of the internal noise and decision bias per subject, which were fit independently using psychometric functions. All models were hierarchically structured and fit to all participants’ trial-wise decisions and confidence ratings together using an Markov chain Monte Carlo (MCMC) approach in STAN^55^ and with the ‘cmdstanr’ package^56^. R-hat values were close to 1 (<1.1) for all parameters. One outlier participant was removed from the hierarchical modelling due to a *w*_*conf*_ parameter that was 6.05 SD from the group mean (or 485.74 SD from the group mean when fit without them) which hence skewed the group-level fit. We analysed the posterior distributions using 89% credible intervals, following the suggestion that these are more stable than 95% intervals for analysing Bayesian posterior distributions^57^. Details of the model implementation in STAN, the model fitting procedure, and the model simplification used can be found in Supplementary Materials.

#### Fitting Internal Noise and Decision Bias

To measure the internal noise (***σ***_prior_ and ***σ***_likelihood_) as well as the decision bias for each participant, we used the approach taken by Lisi et al.^28^, and adapted the scripts available at https://osf.io/w74cn/. We assumed ***σ***_prior_ and ***σ***_likelihood_ to be the same, as we used the same stimuli for leads and targets. We fit four different psychometric functions to participants’ decisions in the control task, as well as the first decision of each main task trial, as these decisions all took place without informative priors (50/50 chance of right vs left). The four psychometric functions were (1) a simple function that included only the internal noise as a free parameter, (2) a function with internal noise as well as decision bias, (3) a function with internal noise as well as a lapse term, and finally (4) a function that included internal noise, decision bias, as well as lapse. The lapse term accounted for the possibility that participants might have made stimulus-independent lapses such as attentional or motor lapses. These four functions were fit and we then used the parameter values retrieved from taking an Akaike-weighted combination of the four estimates. For the modelling analysis, we transformed each participants’ raw coherence values (*coh*) into units of their own internal noise. We additionally transformed their right versus leftward coherence values to take into account their own decision bias. Together, this transformation yielded the following definition of normalized stimulus strength *s*:

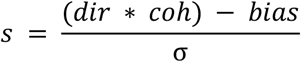

where dir is equal to -1 for leftward stimuli and +1 for rightward stimuli. This transformation allowed us to set the internal noise to 1 in all equations below.

#### Flexible Model

The flexible model included three free parameters per participant - the prior weighting parameter at the decision level (*w*_*choice*_), the prior weighting parameter at the confidence level (*w*_*conf*_), and the confidence bias parameter (*b*). *w*_*choice*_ quantified the relative influence of the prior (compared to the likelihood) in the target decision. This influence of the prior could be captured computationally by shifting the decision criterion. Then, the probability of choosing right (*Φ*_right_) in the target decision was based on the probability that the perceived target stimulus was to the right of the shifted decision criterion. The decision criterion (*θ*) was shifted proportionally to the weighted prior:

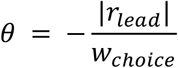

where *r*_*lead*_ is the internal response generated from the lead stimulus and internal noise. This shifting of the decision criterion is computationally equivalent to having a rightward prior equal to the decision confidence on the lead decision (Figure 7), but with the prior variance misestimated according to *w*_*choice*_^28^. The likelihood of a rightward target decision was then computed, exactly as in the work of Lisi et al.^28^, as:

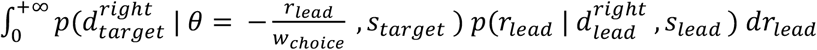

following a rightward lead decision, and as:

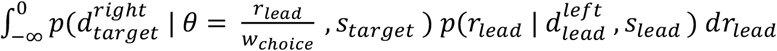

following a leftward lead decision, where *s* represents the stimulus and *d* represents the decision. Because we as experimenters do not have access to the internal signals of the participant, the left term in the integral captures the probability of an internal target signal to the right of the shifted decision criterion – shifted according to the weighted prior signal, which derives from the lead stimulus. The right term weights this by the likelihood of that internal prior signal, given the lead stimulus, and these terms are marginalised across the possible prior signal values.

**Figure 7.**
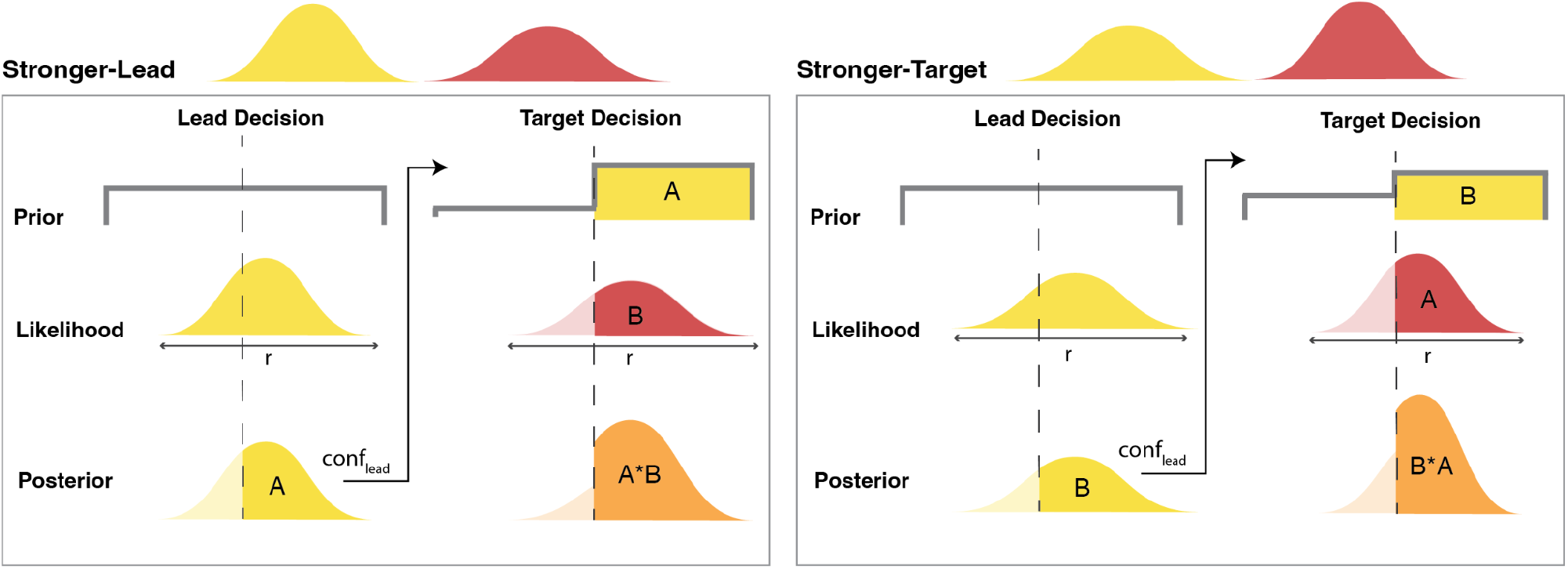
Model Schematic. According to the model, in both conditions, the lead decision occurred under a flat prior. The lead stimulus then generated an internal response with some added internal noise, which formed the likelihood distribution and then the posterior. The area of the posterior on the chosen side of the decision boundary (vertical dashed line) corresponds to the confidence in the lead decision (*conf*_*lead*_), captured by area A (Stronger-Lead condition panel) or B (Stronger-Target condition panel). This confidence in the lead decision then formed the strength of the prior for a rightward target stimulus. The target stimulus generated an internal response that formed the target likelihood, and the prior was then integrated with the target likelihood to give the posterior for the target (orange). This posterior then led to the target decision and confidence rating. The strength of the posterior probability of the winning hypothesis, depicted by the opaque orange area, is based on the combination of the strength of the prior – from the confidence in the lead stimulus – and the strength of the likelihood – from the confidence in the target stimulus. Our two conditions, Stronger-Lead (left) and Stronger-Target (right), simply swapped the strengths of the lead versus target stimulus. But, the confidence in these stimuli (areas A and B) still combine to the same posterior strength (A*B or B*A). Hence, the model predicted equal accuracy and mean confidence between conditions, if the relative weighting of priors and likelihoods was optimal. The weighting parameters, *w*_*choice*_ and *w*_*conf*_, acted by scaling the estimated variance of the lead, effectively scaling *conf*_*lead*_ and capturing the strength of prior for a rightward target that best explains target decisions and confidence ratings, respectively.

Confidence was then modelled as the perceived posterior probability of being correct, combining the prior and likelihood (Figure 7). The relative influence of the prior on confidence was captured by *w*_*conf*_. This weighted rightward prior in confidence (*p(R)*_*conf*_) was equal to the decision confidence from the lead decision, with the variance of the prior misestimated according to *w*_*conf*_ :

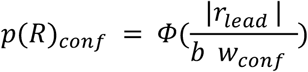

The strength of the likelihood depended on the incoming information from the target stimulus, and was defined as the likelihood of having gotten the internal target signal (*r*_*target*_) if there had been a rightward target stimulus (*R*, or *s*_*target*_>0):

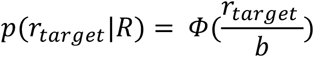

Confidence bias, *b*, captured misestimation of the prior variance as well as an equal misestimation of the likelihood variance, therefore reflecting an overall over- or underconfidence. The posterior combined the prior and likelihood according to Bayes rule, and confidence in a rightward choice was then computed as:

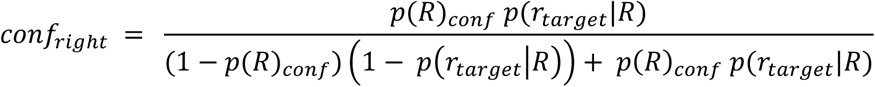

Confidence in a leftward choice was then equal to (1 - *conf*_*right*_). Similarly to *w*_*choice*_, *w*_*conf*_ captured the prior strength that would account for each confidence rating by scaling the variance of the internal prior signal relative to the internal likelihood signal. By implementing these two weighting parameters in the same way, we could then directly compare them. The Flexible model was the only model in which the weighting of the prior information was allowed to differ between discrimination decisions and confidence ratings.

#### Flat Prior Model

The flat prior model captured decisions and confidence in the same way as the Flexible model except that the prior information had no influence on the target decision or confidence, so the lead and target decisions were modelled as independent and confidence was modelled as the decision confidence about only the target stimulus. Computationally this meant forcing the prior for a rightward target stimulus to be equal to 0.5, or an uninformative prior, which was analogous to setting *w*_*choice*_ and *w*_*conf*_ to be infinitely large. The only free parameter in this model was the confidence bias (*b*).

#### Optimal Observer Model

The optimal observer only differed from the Flexible model in that it assumed the prior information to be optimally precision-weighted relative to the likelihood. This meant that *w*_*choice*_ and *w*_*conf*_ were both equal to 1, with only *b* as a free parameter.

#### Equal Model

The equal model was also the same as the Flexible model except that the use of the prior information was assumed to be the same in decisions and confidence, although it could stray from optimal. Computationally, this meant that *w*_*choice*_ and *w*_*conf*_ were forced to be equal to one another, and so only one weighting parameter (*w*) was fit, which was then used as both *w*_*choice*_ and *w*_*conf*_. Again, *b* was still fit to capture an overall confidence bias in this model.

#### Model Comparison

We compared the ability of our four models to account for the behavioural data of the remaining 20 participants after removing the outlier participant. To do this, we performed a 10-fold LOGO-CV, in which we left out 2 participants at a time, fit each model to the remaining 18 participants, and then measured the predictive performance of those fit models for predicting the data of the left-out participants using the ‘loo’ package^58^. The log predictive density for each model for each fold was stored and we then computed the overall expected log pointwise predictive density for each model, and compared them. We considered a model to fit the data better if the magnitude of the difference in expected log pointwise predictive density (elpd_diff) was at least 4, and at least 2 times larger than the standard error of the difference (se_diff)^59^.

## Acknowledgments

We thank Martin Krueck for help with an earlier version of the modelling presented here. MC was supported by the Deutsche Forschungsgemeinschaft (DFG, German Research Foundation) - 337619223 / RTG2386. MC and EF were supported by a Freigeist Fellowship to EF from the Volkswagen Foundation (grant number 91620). MP was supported by a Postdoc. Mobility fellowship from the Swiss National Science Foundation (P400PM_199251). NF has received funding from the European Research Council (ERC) under the European Union’s Horizon 2020 research and innovation program (grant agreement no. 803122). The funders had no role in the conceptualization, design, data collection, analysis, decision to publish, or preparation of the manuscript.

## Data Availability

Raw data is publicly available under https://gitlab.com/MarikaConstant/priors-in-confidence.

## Code Availability

Reproducible analysis scripts and models are publicly available under https://gitlab.com/MarikaConstant/priors-in-confidence.

## Supplementary Materials

### Flexible Model Definition

In the model, the lead stimulus, *s*_*lead*_, generated an internal signal, *r*_*lead*_, by adding normally distributed internal noise, *Ɲ*(0, ***σ***_prior_). The lead decision was made by comparing this internal signal to the decision criterion – signals to the right of the criterion led to a rightward decision, and signals to the left led to a leftward decision. This lead decision then also led to an internal confidence value according to:

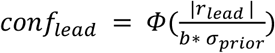

where b is the confidence bias, captured by a general over- or underestimation of the signal variance. The strength of the rightward prior in the target decision was then equal to this *conf*_*lead*_, weighted by the weighting parameter at the decision level, *w*_*choice*_:

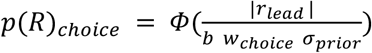

The target stimulus, which was rightward if the lead decision was correct and leftward if not, then generated an internal signal, *r*_*target*_, by adding normally distributed internal noise, *Ɲ*(0, ***σ***_likelihood_). The target decision was based on the posterior probability of a rightward target stimulus, which was determined by the integration of the likelihood and the prior. As shown previously by Lisi et al.^28^, this was computationally equivalent to shifting the target decision criterion and comparing *r*_*target*_ to that shifted decision criterion:

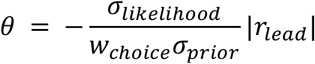

Then, explicit confidence in the target decision was computed as the perceived posterior probability correct, given the evidence and decision. The posterior was again the product of the likelihood and weighted prior, but now the prior was weighted according to the weighting parameter at the confidence level, *w*_*conf*_:

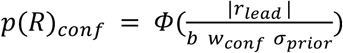

to give:

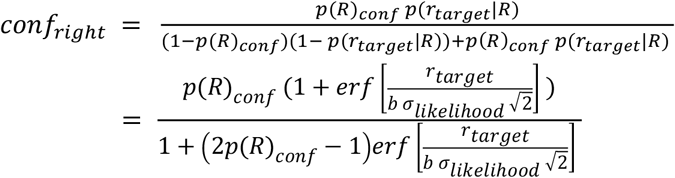

This was the predicted confidence (model_conf) following rightward decisions and *conf*_*left*_ = (1-*conf*_*right*_) was model_conf following leftward decisions.

### Model Fitting

We fit each model using 6000 effective samples across 3 chains, as well as 2000 burn-in samples per chain. All initial parameter values were sampled from a uniform distribution between 0.5 and 1.5. We ensured that all R-hat values were lower than 1.1, indicating good convergence. Lognormal priors were set on the group mean parameters - *w*_*choice*_*_mu, w*_*conf*_*_mu*, and *b_mu*, and subject-wise parameters *w*_*choice*_, *w*_*conf*_ and *b* were modelled as being normally distributed around group mean parameters. Target decisions were modelled as being Bernoulli distributed. Following the procedure used in previous work with a similar modelling approach^60^, we allowed for a small degree of noise (σ=0.0125) between the predicted confidence outlined above (model_conf) and the observed confidence ratings, accounting for some imprecision of participants’ confidence ratings. Posterior predictive checks revealed the model to well capture choice probabilities, as well as differences between conditions in confidence (Figure S1), although confidence was predictably underestimated due to the simplification, explained below.

Priors:

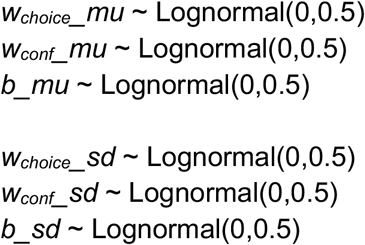

Model:

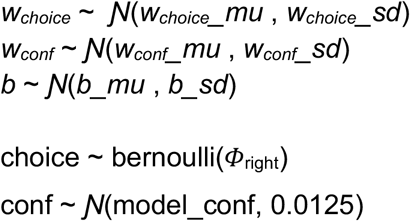

**Figure S1.**
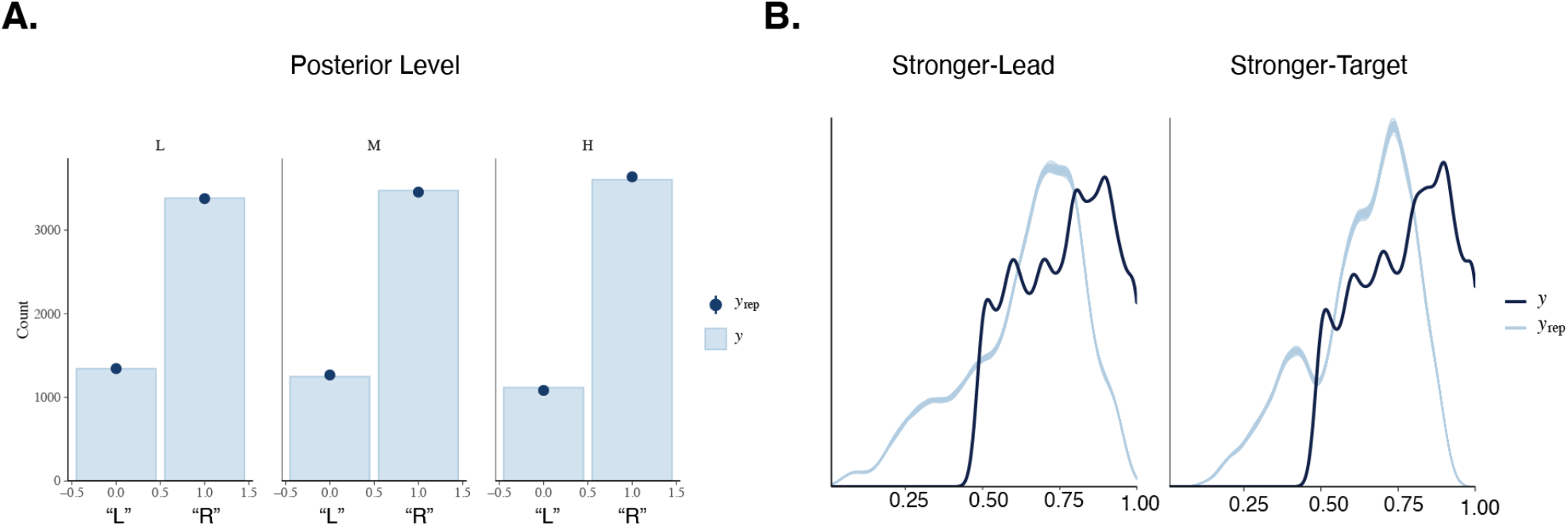
Posterior Predictive Checks. **A. Modelling discrimination decisions**. The light blue bars (y) show the true choice probabilities per posterior level in the data, with rightward target decisions shown at 1, and leftward target decisions shown at 0. The dark blue points (y_rep_) reflect the predicted choice probabilities from the model, generated by sampling from the posterior distribution and simulating target decisions from those samples. **B. Modelling confidence**. In dark blue are the kernel density distributions of confidence in the data per condition (y). In light blue are the predicted kernel density distributions for confidence per condition from the model (y_rep_), generated by sampling from the posterior distribution and simulating target decisions and then confidence from those samples. These are split by the condition to demonstrate that, although the model predictably underestimates confidence, y_rep_ is shifted relative to y similarly in both conditions, therefore allowing us to capture differences in conditions, and this is our primary interest.

### Model Simplification

The *Φ*_right_ and model_conf values were computed as outlined in the main text, however, instead of fitting internal signal values *r*_*lead*_ and *r*_*target*_ on every trial, which would lead to an overparameterized model and complexity issues in the sampling procedure, we simplified the model by fitting confidence based on the external stimulus values. The external stimulus values reflect the mean of the internal signal distribution. We still used the internal noise in the model as otherwise specified. The choice probability was computed by taking the integral across all possible *r*_*lead*_ values, weighted by their likelihood, and hence was not impacted by this simplification. For confidence however, fixing the internal signals at the means meant that the model underestimated confidence in expected ways, particularly following incorrect choices. This primarily impacted the confidence bias parameter *b*, and because this was particularly strong following incorrect trials, *b* was skewed to be inflated in order to increase the value of conf_Incorrect_, and in turn decrease the value of confidence in the correct option (conf_Correct_) which is equal to 1-conf_Incorrect_. Admittedly, this simplification affected not only the bias parameter *b* but also slightly impacted *w*_*conf*_. However we note that it had an effect in a direction directly against our results, as it increased w_conf_. We confirmed this with a parameter recovery analysis in which we simulated data with free internal signals but fit the simplified model, which led to slightly inflated recovery of *w*_*conf*_ (Figure S2). Hence, if we assume participants to also have internal signals, we can expect their true *w*_*conf*_ values to be slightly lower than the values we fit, which is even further in line with our finding that *w*_*conf*_ was lower than *w*_*choice*_.

**Figure S2.**
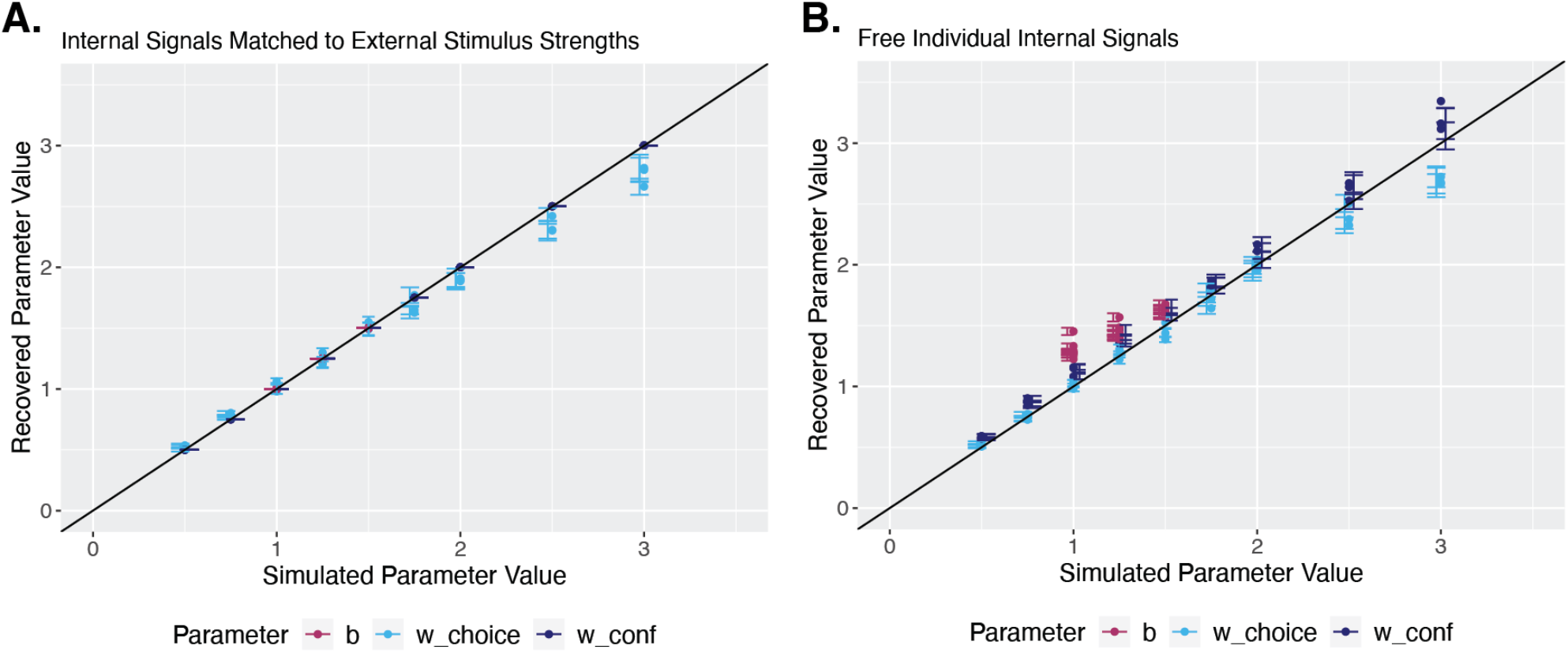
Parameter Recovery Analyses. **A. Fixed internal signals (simplified model)**. Parameter recovery after simulating from the simplified model. The black line indicates perfect parameter recovery. This reveals good parameter recovery of the model when data is simulated from it. Points reflect the mean recovered parameter values across 20 repetitions of simulating data and fitting the model, and error bars reflect SEM across these repetitions. **B. Free internal signals**. Parameter recovery after simulating internal signals but then fitting with the simplified model. This reveals parameters to be reasonably well recovered, despite the simplification. The largest impact is on recovery of *b*, which is inflated relative to the simulated confidence bias. The simplification also leads to a slightly inflated recovery of *w*_*conf*_, but this works directly against our conclusion that *w*_*conf*_ is smaller than *w*_*choice*_, and therefore that the prior is weighted more strongly in confidence than in the decision. Points reflect the mean recovered parameter values across 20 repetitions of simulating data and fitting the model, and error bars reflect SEM across these repetitions.

We also include a model in which we forced any confidence below 0.5 to be set to 0.5, the minimum of the scale, to prevent the skewing of parameters, and that model confirms these results (Figure S3), although there were some convergence challenges when introducing the lower confidence bound.

**Figure S3.**
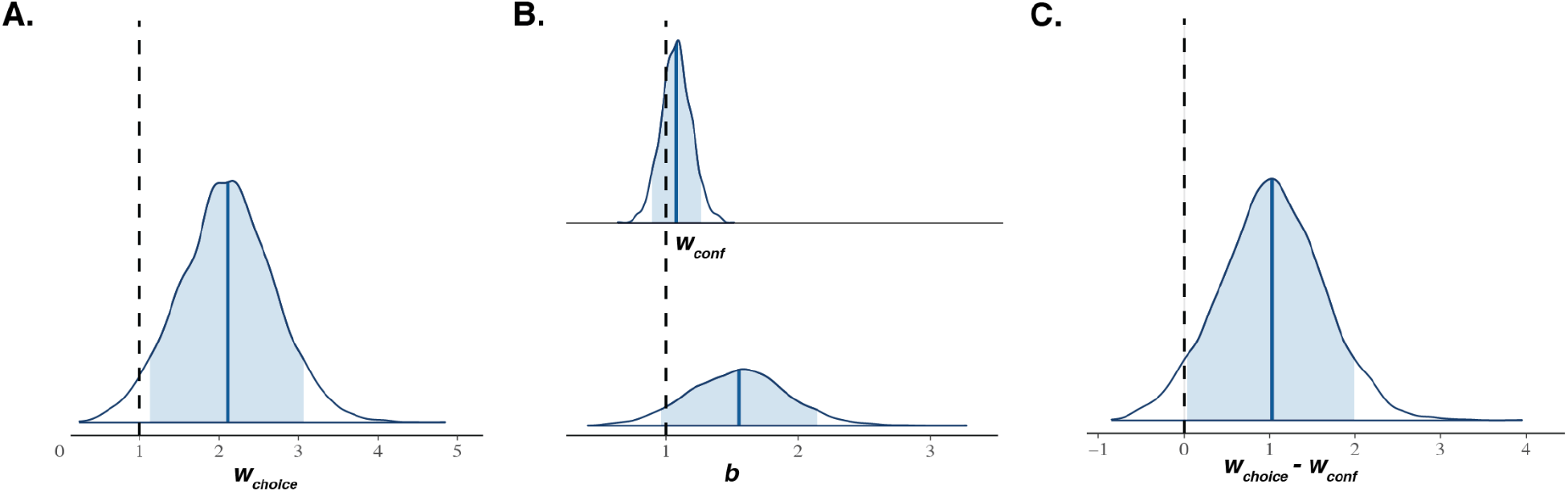
Posterior Group Parameter Distributions for the Bounded Confidence Model. In the bounded confidence model, everything was the same as in the Flexible Model except that any predicted confidence below 0.5 was set to 0.5, which was the minimum of the scale provided to participants. This prevented skewing of parameters due to the model simplification, described above. However, there were some convergence issues due to the discontinuity this introduced. All group parameters converged (R-hat < 1.1) but four individuals within the model had parameters that did not converge. We were, however, able to fit the model individually, and then it converged for all except one participant, and led to similar estimates as in the hierarchical fit. **A. Posterior distribution for *w***_***choice***_. The posterior distribution for the group mean parameter of the weighting of prior information in the decision, *w*_*choice*_. The blue shaded region shows the 89% credible interval and the vertical black dashed line reflects optimal weighting of the prior in the decision (*w*_*choice*_ = 1). **B. Posterior distribution for *w***_***conf***_ **and *b***. The top posterior distribution is for the group mean parameter of *w*_*conf*_. The lower posterior distribution is for the group mean parameter of *b*. The blue shaded regions show the 89% credible intervals and the vertical black dashed line corresponds to the parameter values of an optimal observer. Note that the estimates for both of these parameters are lower here in comparison to the unbounded version of the model, as this helps reduce the skew due to the model simplification. **C. Posterior Group Difference Distribution of *w***_***choice***_ ***- w***_***conf***_. The posterior distribution for the difference in the group mean parameters *w*_*choice*_ and *w*_*conf*_. The blue shaded region shows the 89% credible interval and the vertical black dashed line reflects no difference in the two parameters (*w*_*choice*_ - *w*_*conf*_ = 0). 0 is excluded from the 89% credible interval, suggesting *w*_*choice*_ and *w*_*conf*_ to be credibly different from one another.

### Model Recovery

In order to ensure that our models were adequately distinguishable from one another, we performed a model recovery analysis in which we simulated data from parameters that were associated with the different models, and checked whether the correct models were recovered. We simulated data from each of the simpler models, as well as two parameter combinations of the Flexible Model, one case with *w*_*choice*_ greater than *w*_*conf*_, and one with that reversed, and then compared the model fits in each case using the PSIS-LOO CV approach^58^. This was repeated 10 times for every model/parameter combination. We considered a model to better predict the data if the ELPD was higher by at least 4, and with a magnitude of difference that was at least 2 times the standard error of the difference. We expected that, when fitting data simulated from the Equal Model, both the Flexible and Equal Model would be able to predict the data, since the Flexible Model is able to give equal *w*_*choice*_ and *w*_*conf*_ parameters as well. When data were simulated from the Optimal Model, we expected the Flexible, Equal, and Optimal Models to all predict the data equally well, since all three models can account for *w*_*choice*_ and *w*_*conf*_ values of 1. The most critical test for us was that the Flexible Model was only a *superior* model fit when the data came from that model. This was confirmed with the model recovery analysis, with the Flexible Model winning only when data was simulated from the Flexible Model, in either parameter combination (Figure S4). Also in line with our expectations, data from the Equal Model was best predicted by either the Equal Model or Flexible Model, which were not distinguishable; data from the Optimal Model was best predicted by either the Optimal Model, Equal Model, or Flexible Model, which were not distinguishable; and data from the Flat Prior Model was best predicted by the Flat Prior Model (Figure S4). This confirmed that the Flexible Model was distinguishable from the simpler models, in that it would only win the comparison when data was truly simulated from that model, and hence *w*_*choice*_ and *w*_*conf*_ were different from optimal, different from each other, and different from infinite (flat prior).

**Figure S4.**
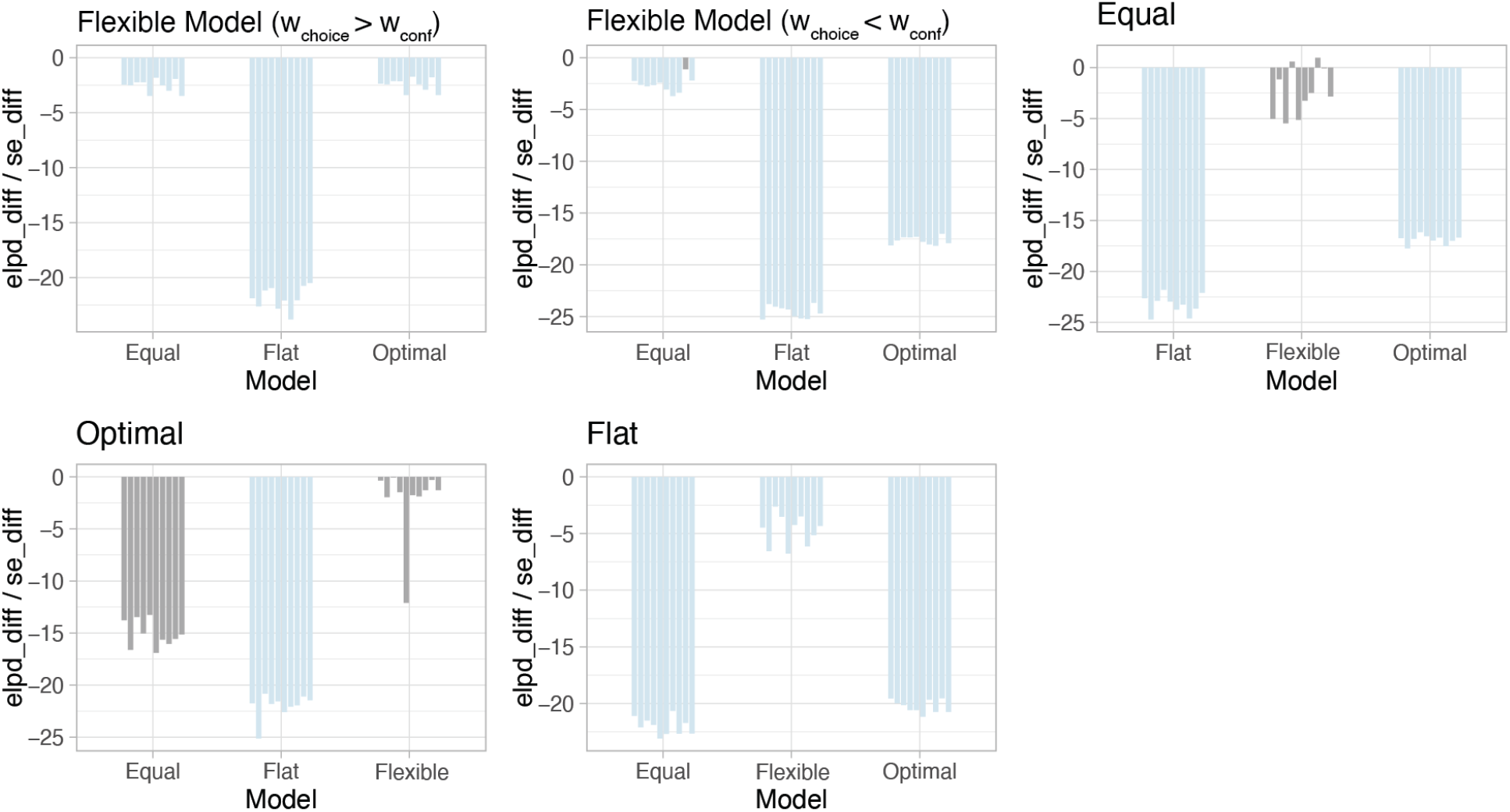
Model Recovery Analysis. To test model recovery, we simulated data from each model and then compared the predictive performance of all models on that data. We consider a model to perform better when the ELPD is higher by at least 4, and when the magnitude of the difference is at least 2 times the standard error of the difference. Bars indicate the difference in ELPD divided by the SE of the difference, always in comparison to the true model, labelled above each facet. Negative values suggest the model to have a lower ELPD than the true model, but bars are coloured light blue if this difference meets the other criteria, namely having a magnitude of at least 4 and at least 2 times the standard error of the difference. Therefore, negative light blue bars indicate a significantly worse predictive performance, compared to the true model. Bars are coloured grey if these criteria are not met and hence the models are not distinguishable. We simulated data from the Flexible Model using two different parameter combinations – one in which *w*_*choice*_ was larger than *w*_*conf*_ (*w*_*choice*_ = 2, *w*_*conf*_ = 1) and one in which this was reversed (*w*_*choice*_ = 1, *w*_*conf*_ = 2). For the Equal Model, we simulated data with the weighting parameter *w* set to 2 (for both decisions and confidence). For the Optimal Model, the weighting of prior information was optimal (1 in both decisions and confidence). For the Flat Prior Model, decisions and confidence were simulated without any use of prior information (equivalent to *w*_*choice*_ and *w*_*conf*_ being infinitely large). Importantly, the only time in which the Flexible Model was conclusively superior in terms of predictive performance was when data were simulated from that model, and hence when *w*_*conf*_ and *w*_*choice*_ were really different than optimal, different than each other, and not infinitely large, suggesting the models to be distinguishable in the required way.

### Experiment 2

**Figure S5.**
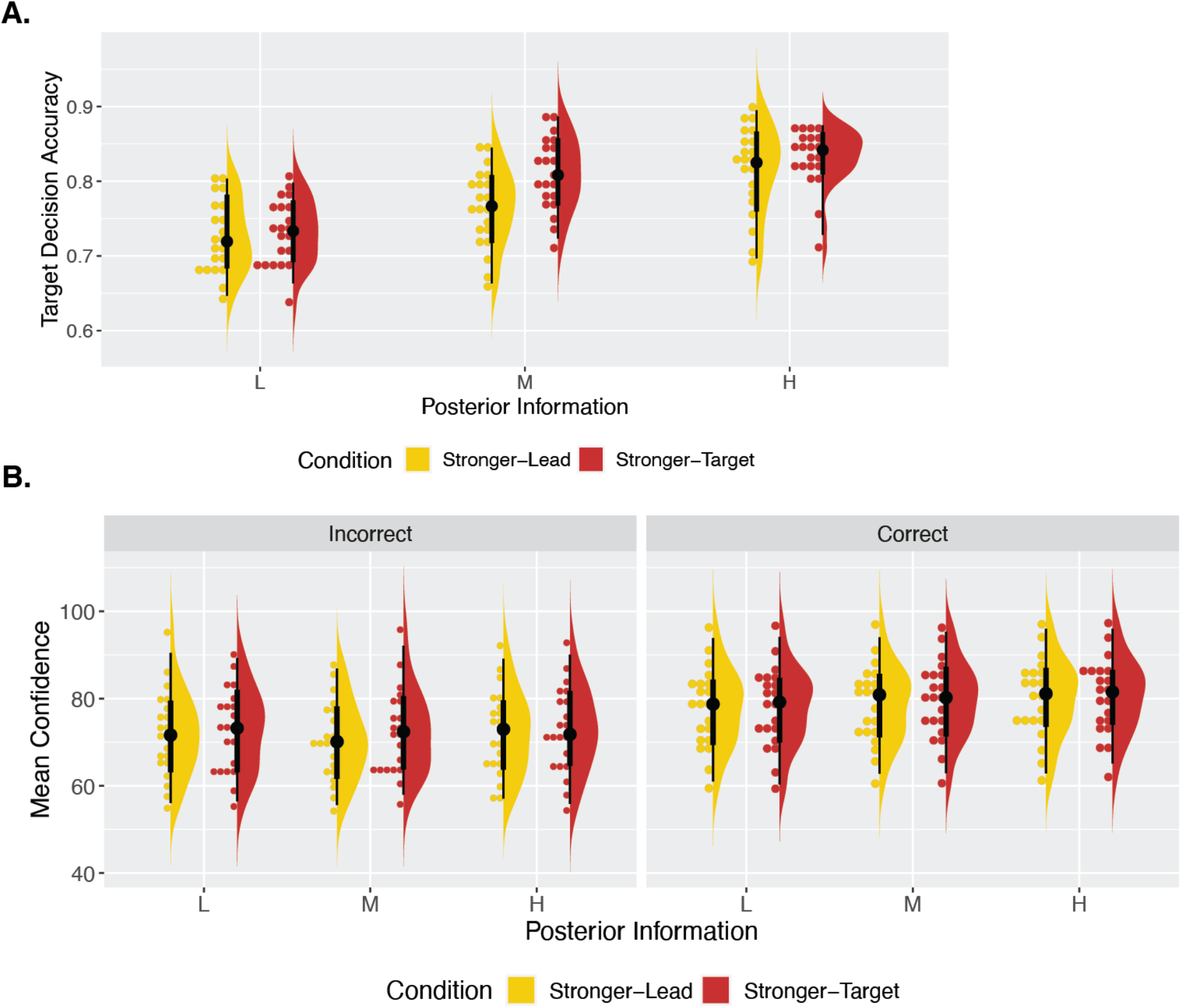
Experiment 2 Behavioural Results. **A. Effect of condition on accuracy**. We found a significant interaction between the effects of condition (which stimulus was stronger) and posterior level on target decision accuracy. Decision accuracy was higher in the Stronger-Target condition and this effect was the strongest at the medium posterior level, suggesting participants to underweight prior information in their decisions, as in Experiment 1. In the raincloud plots, the right-half, split violin plots show the probability density, and vertical black lines show the median, IQR, hinges showing the first and third quartiles, and vertical whiskers showing +/- 1.5IQR. The binned dotplots on the left half show each individual subject as a point. **B. Effect of condition on confidence**. We found a significant effect of condition on confidence, with higher confidence in the Stronger-Target condition.

## References

1. Kersten, D., Mamassian, P. & Yuille, A. Object Perception as Bayesian Inference. Annu. Rev. Psychol. 55, 271–304 (2004).

2. Daw, N. D., Gershman, S. J., Seymour, B., Dayan, P. & Dolan, R. J. Model-Based Influences on Humans’ Choices and Striatal Prediction Errors. Neuron 69, 1204–1215 (2011).

3. Friston, K. et al. Active inference and learning. Neurosci. Biobehav. Rev. 68, 862–879 (2016).

4. Ma, W. J. & Jazayeri, M. Neural coding of uncertainty and probability. Annu. Rev. Neurosci. 37, 205–220 (2014).

5. McNamara, J. M., Green, R. F. & Olsson, O. Bayes’ theorem and its applications in animal behaviour. Oikos 112, 243–251 (2006).

6. Wiese, W. & Metzinger, T. Vanilla PP for Philosophers: A Primer on Predictive Processing. in Philosophy and Predictive Processing (eds. Metzinger, T. & Wiese, W.) (2017).

7. Zellner, A. Bayesian and non-Bayesian approaches to statistical inference and decision-making. J. Comput. Appl. Math. 64, 3–10 (1995).

8. Aitchison, L., Bang, D., Bahrami, B. & Latham, P. E. Doubly Bayesian Analysis of Confidence in Perceptual Decision-Making. PLOS Comput. Biol. 11, e1004519 (2015).

9. Fleming, S. M. & Daw, N. D. Self-Evaluation of Decision-Making: A General Bayesian Framework for Metacognitive Computation. Psychol. Rev. 124, 91–114 (2017).

10. Kepecs, A. & Mainen, Z. F. A computational framework for the study of confidence in humans and animals. Philos. Trans. R. Soc. B Biol. Sci. 367, 1322–1337 (2012).

11. Kiani, R. & Shadlen, M. N. Representation of Confidence Associated with a Decision by Neurons in the Parietal Cortex. Science 324, 759–764 (2009).

12. Meyniel, F., Sigman, M. & Mainen, Z. F. Confidence as Bayesian Probability: From Neural Origins to Behavior. Neuron 88, 78–92 (2015).

13. Pouget, A., Drugowitsch, J. & Kepecs, A. Confidence and certainty: distinct probabilistic quantities for different goals. Nat. Neurosci. 19, 366–374 (2016).

14. Sanders, J. I., Hangya, B. & Kepecs, A. Signatures of a Statistical Computation in the Human Sense of Confidence. Neuron 90, 499–506 (2016).

15. Marcus, G. F. & Davis, E. How robust are probabilistic models of higher-level cognition? Psychol. Sci. 24, 2351–2360 (2013).

16. Douven, I. & Schupbach, J. N. Probabilistic alternatives to Bayesianism: the case of explanationism. Front. Psychol. 6, (2015).

17. Peters, M. A. K. et al. Perceptual confidence neglects decision-incongruent evidence in the brain. Nat. Hum. Behav. 1, (2017).

18. Navajas, J. et al. The idiosyncratic nature of confidence. Nat. Hum. Behav. 1, 810–818 (2017).

19. Rausch, M., Hellmann, S. & Zehetleitner, M. Confidence in masked orientation judgments is informed by both evidence and visibility. Atten. Percept. Psychophys. 80, 134–154 (2018).

20. Pleskac, T. J. & Busemeyer, J. R. Two-stage dynamic signal detection: A theory of choice, decision time, and confidence. Psychol. Rev. 117, 864–901 (2010).

21. van den Berg, R. et al. A common mechanism underlies changes of mind about decisions and confidence. eLife 5, e12192 (2016).

22. Charles, L. & Yeung, N. Dynamic Sources of Evidence Supporting Confidence Judgments and Error Detection. J. Exp. Psychol. Hum. Percept. Perform. 45, (2018).

23. Murphy, P. R., Robertson, I. H., Harty, S. & O’Connell, R. G. Neural evidence accumulation persists after choice to inform metacognitive judgments. eLife 4, e11946 (2015).

24. Balsdon, T., Wyart, V. & Mamassian, P. Confidence controls perceptual evidence accumulation. Nat. Commun. 11, 1753 (2020).

25. Balsdon, T., Mamassian, P. & Wyart, V. Separable neural signatures of confidence during perceptual decisions. eLife 10, e68491 (2021).

26. Locke, S. M., Gaffin-Cahn, E., Hosseinizaveh, N., Mamassian, P. & Landy, M. S. Priors and payoffs in confidence judgments. Atten. Percept. Psychophys. 82, 3158–3175 (2020).

27. Sherman, M. T., Seth, A. K., Barrett, A. B. & Kanai, R. Prior expectations facilitate metacognition for perceptual decision. Conscious. Cogn. 35, 53–65 (2015).

28. Lisi, M., Mongillo, G., Milne, G., Dekker, T. & Gorea, A. Discrete confidence levels revealed by sequential decisions. Nat. Hum. Behav. 5, 273–280 (2021).

29. Pereira, M. et al. Evidence accumulation relates to perceptual consciousness and monitoring. Nat. Commun. 12, 3261 (2021).

30. Pereira, M., Perrin, D. & Faivre, N. A leaky evidence accumulation process for perceptual experience. Trends Cogn. Sci. 26, 451–461 (2022).

31. Cicchini, G. M., Benedetto, A. & Burr, D. C. Perceptual history propagates down to early levels of sensory analysis. Curr. Biol. 31, 1245-1250.e2 (2021).

32. de Lange, F. P., Heilbron, M. & Kok, P. How Do Expectations Shape Perception? Trends Cogn. Sci. 22, 764–779 (2018).

33. Teufel, C. & Fletcher, P. C. Forms of prediction in the nervous system. Nat. Rev. Neurosci. 21, 231–242 (2020).

34. Olawole-Scott, H. & Yon, D. Expectations about precision bias metacognition and awareness. Preprint at https://doi.org/10.31234/osf.io/um2wx (2022).

35. Van Marcke, H., Denmat, P. L., Verguts, T. & Desender, K. Manipulating prior beliefs causally induces under- and overconfidence. http://biorxiv.org/lookup/doi/10.1101/2022.03.01.482511 (2022) doi:10.1101/2022.03.01.482511.

36. Rollwage, M. et al. Confidence drives a neural confirmation bias. Nat. Commun. 11, 2634 (2020).

37. Stocker, A. A. & Simoncelli, E. P. A Bayesian Model of Conditioned Perception. Adv. Neural Inf. Process. Syst. 2007, 1409–1416 (2007).

38. Caziot, B. & Mamassian, P. Perceptual confidence judgments reflect self-consistency. J. Vis. 21, 8 (2021).

39. Festinger, L. A Theory of Cognitive Dissonance. (Stanford University Press, 1957).

40. Pellicano, E. & Burr, D. When the world becomes ‘too real’: a Bayesian explanation of autistic perception. Trends Cogn. Sci. 16, 504–510 (2012).

41. Skewes, J. C., Jegindø, E.-M. & Gebauer, L. Perceptual inference and autistic traits. Autism 19, 301–307 (2015).

42. Van de Cruys, S. et al. Precise minds in uncertain worlds: Predictive coding in autism. Psychol. Rev. 121, 649–675 (2014).

43. Adams, R., Stephan, K., Brown, H., Frith, C. & Friston, K. The Computational Anatomy of Psychosis. Front. Psychiatry 4, (2013).

44. Corlett, P. R., Frith, C. D. & Fletcher, P. C. From drugs to deprivation: a Bayesian framework for understanding models of psychosis. Psychopharmacology (Berl.) 206, 515–530 (2009).

45. Fletcher, P. C. & Frith, C. D. Perceiving is believing: a Bayesian approach to explaining the positive symptoms of schizophrenia. Nat. Rev. Neurosci. 10, 48–58 (2009).

46. Sterzer, P. et al. The Predictive Coding Account of Psychosis. Biol. Psychiatry 84, 634–643 (2018).

47. Lange, K., Kühn, S. & Filevich, E. “Just Another Tool for Online Studies” (JATOS): An Easy Solution for Setup and Management of Web Servers Supporting Online Studies. PLOS ONE 10, e0130834 (2015).

48. Rajananda, S., Lau, H. & Odegaard, B. A Random-Dot Kinematogram for Web-Based Vision Research. J. Open Res. Softw. 6, 6 (2018).

49. Bates, D., Mächler, M., Bolker, B. & Walker, S. Fitting Linear Mixed-Effects Models Using lme4. J. Stat. Softw. 67, 1–48 (2015).

50. R Core Team. R: A language and environment for statistical computing. (2021).

51. Barr, D. J., Levy, R., Scheepers, C. & Tily, H. J. Random effects structure for confirmatory hypothesis testing: Keep it maximal. J. Mem. Lang. 68, 255–278 (2013).

52. Makowski, D., Ben-Shachar, M. S. & Lüdecke, D. bayestestR: Describing Effects and their Uncertainty, Existence and Significance within the Bayesian Framework. J. Open Source Softw. 4, 1541 (2019).

53. Bürkner, P.-C. brms: An R Package for Bayesian Multilevel Models Using Stan. J. Stat. Softw. 80, 1–28 (2017).

54. Maniscalco, B. & Lau, H. A signal detection theoretic approach for estimating metacognitive sensitivity from confidence ratings. Conscious. Cogn. 21, 422–430 (2012).

55. Stan Development Team. Stan Development Team. 2022. Stan Modeling Language Users Guide and Reference Manual, 2.30. https://mc-stan.org. (2022).

56. Gabry, J. & Češnovar, R. R Interface to CmdStan. https://mc-stan.org/cmdstanr/ (2022).

57. Kruschke, J. Doing Bayesian Data Analysis: A Tutorial with R, JAGS, and Stan. (Academic Press, 2014).

58. Vehtari, A. et al. loo: Efficient leave-one-out cross validation and WAIC for Bayesian models. (2022).

59. Sivula, T., Magnusson, M., Matamoros, A. A. & Vehtari, A. Uncertainty in Bayesian Leave-One-Out Cross-Validation Based Model Comparison. Preprint at https://doi.org/10.48550/arXiv.2008.10296 (2022).

60. Fleming, S. M., van der Putten, E. J. & Daw, N. D. Neural mediators of changes of mind about perceptual decisions. Nat. Neurosci. 21, 617–624 (2018).

